# Using Single Protein/Ligand Binding Models to Predict Active Ligands for Unseen Proteins

**DOI:** 10.1101/2020.08.02.233155

**Authors:** Vikram Sundar, Lucy Colwell

## Abstract

Machine learning models that predict which small molecule ligands bind a single protein target report high levels of accuracy for held-out test data. An important challenge is to extrapolate and make accurate predictions for new protein targets. Improvements in drug-target interaction (DTI) models that address this challenge would have significant impact on drug discovery by eliminating the need for high-throughput screening experiments against new protein targets. Here we propose a data augmentation strategy that addresses this challenge to enable accurate prediction in cases where no experimental data is available. To proceed, we first build single protein-ligand binding models and use these models to predict whether additional ligands bind to each protein. We then use these predictions to augment the experimental data, train standard DTI models, and predict interactions between unseen test proteins and ligands. This approach achieves Area Under the Receiver Operator Characteristic (AUC) > 0.9 consistently on test sets consisting exclusively of proteins and ligands for which the model is given no experimental data. We verify that performance improvements extend to held-out test proteins distant from the training set. Our data augmentation framework can be applied to any DTI model, and enhances performance on a range of simple models.

## Introduction

Identifying ligands that interact with a given protein target, and predicting additional protein targets that a candidate drug binds to are crucial requirements in drug discovery. Experimental methods such as high-throughput screening are time-consuming and costly, while physics-based methods are computationally expensive and can be inaccurate (1–4). The emergence of large datasets describing experimentally measured interactions between protein targets and small molecule ligands enables data-driven approaches to be applied to this problem (5). In recent years, a variety of machine learning approaches have been developed to identify active ligands for single protein targets using training data from screening experiments (6, 7). These approaches report outstanding *in silico* success on benchmark datasets (8–15). However, they rely on the existence of experimental screening data that identifies active and ideally also inactive ligands for each protein target, which is costly and time-consuming to obtain.

Models that predict global drug-target interactions (DTI) remove this bottleneck by exploiting the physio-chemical information shared across different protein-ligand binding interactions, to simultaneously predict interactions between multiple protein targets and multiple candidate ligands or drugs (16–20). DTI models use the experimental data available for some subset of interactions (Fig. 1A), together with information-rich featurizations of the proteins and ligands, to predict interactions between test proteins and ligands. The ultimate goal is to build models that accurately predict interactions where no experimental data is available, enabling, for example, prediction of active ligands for protein targets with no prior screening data (16, 19). If successful, these models would be particularly useful in the prediction of off-target effects, and would enable better understanding of drug polypharmacology.

**Fig. 1.**
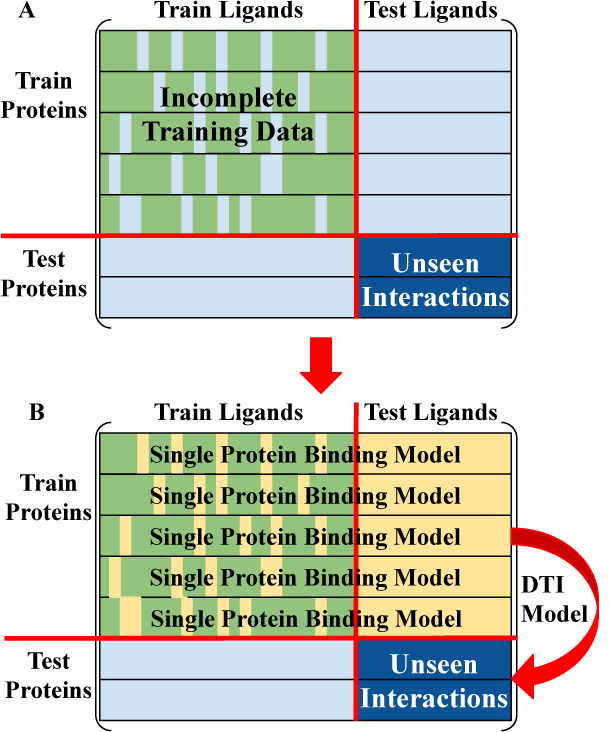
(a) The hardest DTI challenge involves building models that predict interactions between proteins and ligands for which no experimental data is available, which requires generalization in both protein and ligand space. (b) We first use the available data to build a single-target binding model for each training protein, and then use the resulting model to predict interactions between that protein and all ligands in the dataset, augmenting the experimental data (yellow fill). Standard DTI models then generalise from the augmented data and predict interactions between the test proteins and ligands (dark blue).

However, performance analyses on standard benchmark datasets show that DTI methods can struggle to make accurate prediction for interactions between protein targets and unseen ligands (19). This is particularly surprising, since single protein models successfully solve this problem without including data for other protein targets (5). Many deep-learning-based DTI models have been proposed, but they typically rely on large numbers of experimentally determined active ligands for each protein. Performance of these approaches at generalizing to proteins or ligands without experimental data is poor (21–32). Moreover, methods for DTI prediction have largely been tested using a pair splitting methodology, where the dataset of (protein, ligand) pairs is split into train, validation, and test sets (16, 19, 24–26, 32– 42). This approach does not determine whether models can generalize to unseen protein targets, since the test set contains protein targets for which many active ligands are seen during training.

The most difficult and most interesting test case of generalising in both protein and ligand space simultaneously (Fig. 1, dark blue) has rarely been tried in the literature (19); when it is tested, models often perform poorly (19, 21, 29, 31, 43) though in some cases this is partially accounted for by the difficulty of the dataset (29). A small number of papers have used deep learning to explicitly tackle the problem of making predictions for targets with no training data, reporting lower performance than models tested via pair splitting (21, 23, 31). In this paper, we develop a data augmentation framework to tackle this most challenging case and accurately predict interactions in cases where neither the ligands nor the protein targets were seen during training. Specifically, we first use highly accurate single protein/ligand binding models to generalise and make predictions for unseen ligands for each protein, then use these predicted interactions to augment the training data for DTI models that generalise to unseen proteins. Using simple DTI models, we demonstrate that the ensuing performance is comparable or better than that of state-of-the-art deep learning-based DTI models (31). Most importantly, our method makes it possible to tackle the most challenging case involving simultaneous generalisation to both unseen drugs and unseen targets with remarkable accuracy.

## Results

To motivate our approach, we first consider the simpler problem of predicting whether unseen small molecule ligands bind to protein targets with some experimental binding data, for which single protein-ligand binding models have achieved outstanding reported success. We reasoned that these models should provide a baseline for the performance of DTI models at this problem, since the DTI models capture target similarity and so can transfer information between protein targets. However, Figure S1 shows that single protein-ligand binding models outperform DTI models despite having access to less information. It is certainly true that the high performance of single protein/ligand binding models may reflect similarities between train and test ligands, or other forms of dataset bias (44–46). Regardless, we find that DTI models are unable to replicate this performance.

This observation suggests that the accurate single protein models could be leveraged for data augmentation, to provide additional information that helps a DTI model generalise to unseen protein targets. To test this hypothesis, we construct an updated version of the benchmark dataset from (36) that contains a hundredfold more ligands for each protein class. We build single protein/ligand binding models for each protein target in the train set, and use these models to predict interactions between the corresponding protein and all other ligands in the dataset (see Figure 1b). We next train a DTI model using the experimentally validated data augmented by the predicted interactions, and test whether it can generalise to make accurate predictions for unseen proteins. Since we have already made predictions for all ligands in the dataset, we use the DTI method exclusively to generalise in protein space to interactions involving unseen protein targets, avoiding the expensive ligand similarity matrix calculation. We compare performance to that of the DTI model trained using only the experimentally validated interactions. To explicitly test whether data augmentation enables DTI models to generalise in both protein and ligand space simultaneously, we hold out sets of protein targets and small molecule ligands from training, and measure DTI model performance for interactions with no experimental data for either the protein or the ligand (dark blue, Figure 1).

Similar approaches have been applied to related problems; specifically, previous work has found that single proteinligand binding models like random forest can improve accuracy of protein family regression models (47). This work did not incorporate information about the protein targets and therefore required data for every target; in contrast, our work with data augmentation on DTI models can be used even for proteins with no prior experimental data (47). Weighted nearest neighbor was previously used as a data augmentation method in combination with a new DTI model, though its contribution to performance was not evaluated (40).

Figure 2 reports the Area under the Receiver Operator Characteristic (AUC) for three DTI models both before and after data augmentation, across held-out test proteins and held-out test ligands from five data sets (updated from (36), see methods). Results for additional DTI models are shown in Figure S2. All results show that augmenting the training data leads to statistically significant improvements (*p* < 0.01 level). In general, Figure 2 shows greater improvement when predictions from logistic regression models are used for data augmentation. Figures S2-S10 have further analysis of these results, indicating that they hold for other similarity-based DTI methods, for easier prediction tasks involving interactions between unseen ligands and known targets and between unseen targets and known ligands, and also on a trial-by-trial basis. When the augmented training data is used, even baseline models like weighted nearest neighbours (weighted-NN) can achieve high accuracy, with AUCs exceeding 0.85 as shown in Figure S2. This suggests that data augmentation allows even simple DTI models to generalise simultaneously in both protein and ligand space. Many of the more complex similarity-based DTI models demonstrate very similar levels of improved performance following data augmentation. Moreover, while we did not record exact runtimes, our observations indicate that data augmentation significantly reduces both runtime and memory costs since the ligand similarity matrix does not need to be computed and stored, which can take hundreds of gigabytes and several hours of computer time on a single core. Figure S11 demonstrates that incorporating the ligand similarity matrix into the DTI model post-augmentation is detrimental to model performance. Overall, our results suggest that using single protein/ligand binding models to label additional training data points has the potential to relieve a crucial bottleneck in the ability of DTI models to generalise to new protein targets.

**Fig. 2.**
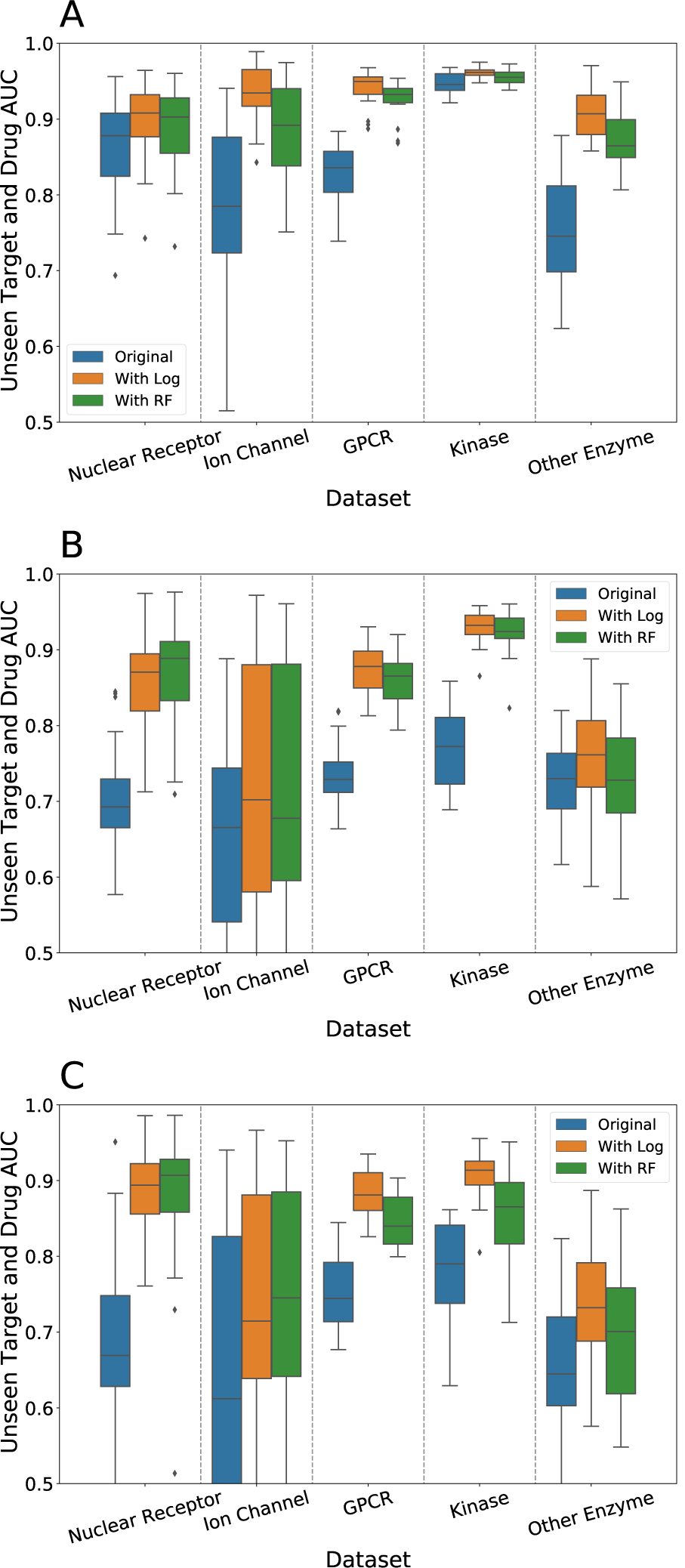
Effect of Data Augmentation on Unseen Target and Drug AUCs for the DTI Models: (a) Random Forest trained using counts of amino acids in the target sequence (RF One-Hot) (b) Collaborative Matrix Factorization (CMF) (38), and (c) Weighted Graph-Regularized Matrix Factorization (WGRMF, see SI Methods for details) (39). We see consistent improvement in all models due to data augmentation using simple single protein/ligand binding models, regardless of dataset size or sparsity and the specific DTI model used. In particular, after augmentation RF one-hot consistently attains AUCs above 0.9, suggesting that it can simultaneously generalise in protein and ligand space. We typically see greater improvement using logistic regression models for data augmentation rather than random forest, but both methods are effective.

### Performance Analysis

For the proposed data augmentation approach to be generally useful it is important to understand how model performance depends on the amount and diversity of available training data. First, we examine how performance depends on the distance of a test protein target from the proteins contained in the training data. For each held-out test protein, we measure the similarity to each training protein using pHMMer (48), and report the normalized bit score of the most similar match (see methods). Figure 3a-c reports the improvement in the DTI model AUC for each test target obtained through data augmentation, as a function of its similarity to the training set. The raw AUC values are reported in Figure 3d-f; Figure S15 depicts the distribution of test target similarities to the training set. We note that as a result of data augmentation, the simple random forest DTI model (RF One-Hot), where target similarity is computed using amino acids counts (see methods) performs highly on protein targets that are highly dissimilar to the training set across the GPCR, Kinase and Other Enzyme datasets. Furthermore, Figure S14 shows that the performance improvement does not depend strongly on the number of known active ligands per protein target.

**Fig. 3.**
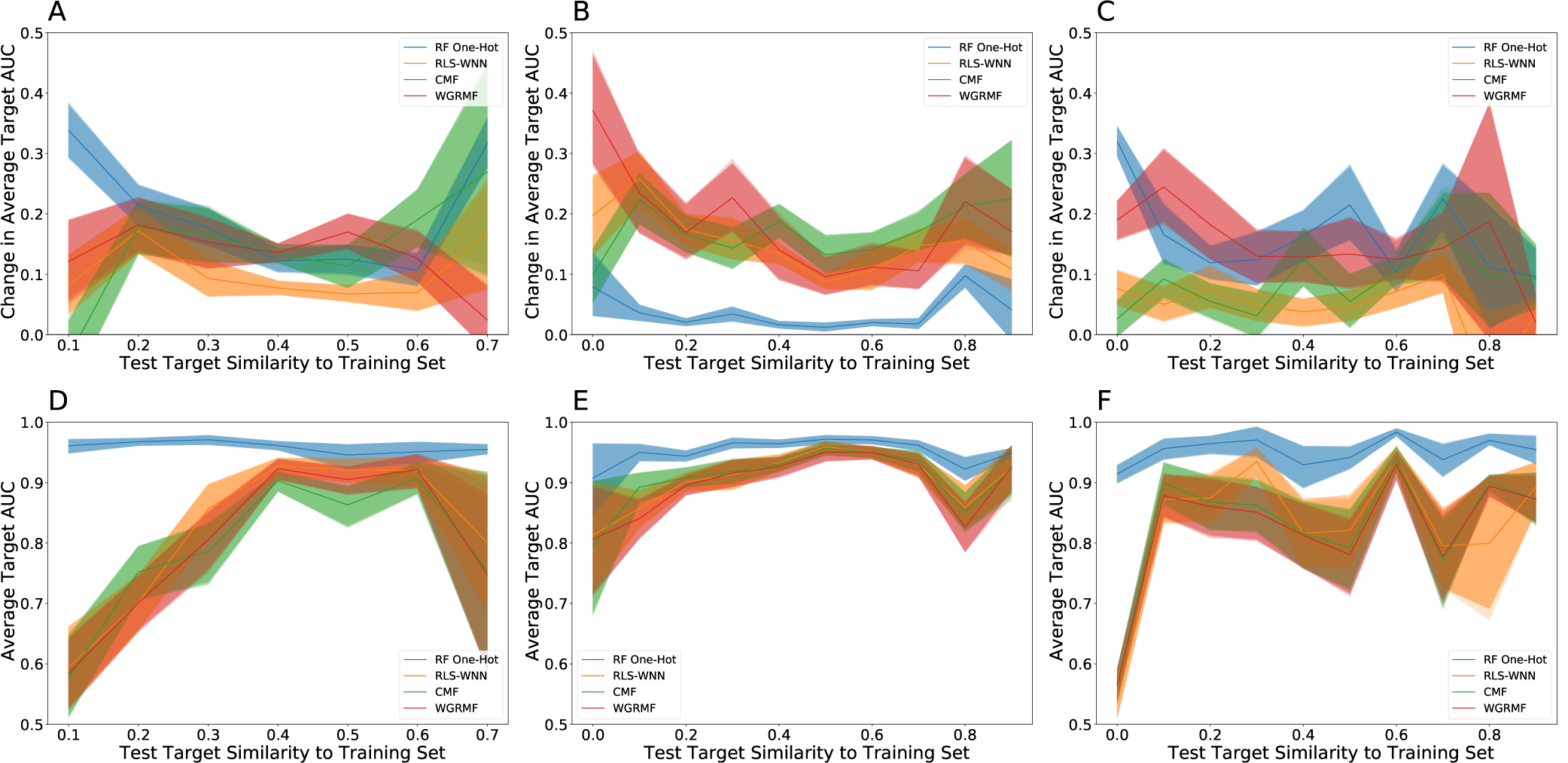
(A-C) Change in Target AUC and (D-F) Target AUC as a Function of Similarity to Training Set on Datasets with More than 50 Protein Targets. (A, D) are the GPCR dataset, (B, E) the Kinase dataset, and (C, F) the Other Enzyme dataset. Our method is effective across test proteins regardless of their similarity to the initial training set. The error bars for this graph were generated by a bootstrap estimate after binning the test target distances into bins of width 0.05.

We can similarly examine how performance depends on the distance of a test ligand to the nearest ligand in the training data, where distance is measured using Tanimoto similarity. Figure 4a-c reports the improvement in the DTI model AUC obtained through data augmentation, as a function of test ligand similarity to the training set. The raw AUC values are reported in Figure 4; Figure S16 depicts the distribution of test ligand similarities to the training set. All models show a decrease in performance as distance from the training set increases, but our data augmentation method is generally effective across a wide range of distances.

**Fig. 4.**
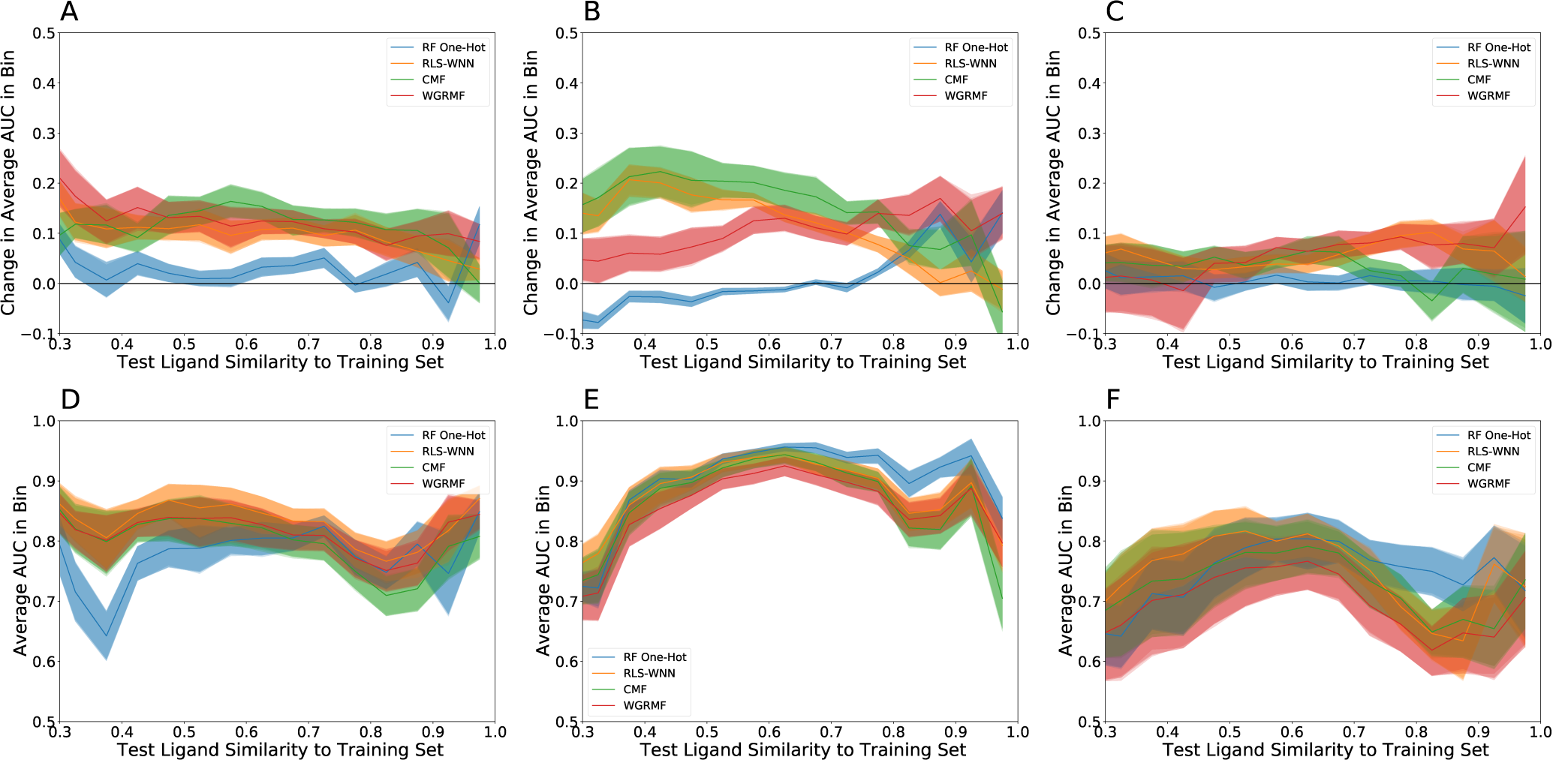
(A-C) Change in AUC and (D-F) Average AUC in Ligand Bin as a Function of Ligand Similarity to Training Set on Datasets with More than 50 Protein Targets. (A, D) are the GPCR dataset, (B, E) the Kinase dataset, and (C, F) the Other Enzyme dataset. Ligands are binned based on distance from the training set in bins of width 0.05. Our method is effective across test ligands regardless of their similarity to the initial training set. The error bars for this graph were generated by a bootstrap estimate.

Next, we examine performance as a function of the diversity of protein targets included in the training set. We first fix a held-out test set of target proteins for the GPCR, kinase and ion channel datasets, using the other proteins in each set as training data. For each dataset, we then remove all data involving the 5% of train protein targets that are on average most dissimilar from the other train proteins, making each training set less diverse in the most rapid order. We repeat for 10 steps to yield 10 train sets of varying diversity for each dataset. We measure the diversity of each training set using the average nearest-neighbor similarity across the protein targets it contains. In general, a less diverse training set has less information, which makes it potentially more difficult to generalise in protein space. However, Figure 5 suggests that the improvement obtained by our data augmentation approach decreases only slightly as the training set becomes less diverse. The data augmented DTI models are consistently effective across a wide range of training set diversities. For comparison, Figure S13 shows similar results where we reduce the size of the training set randomly instead of in the order that decreases diversity most quickly.

**Fig. 5.**
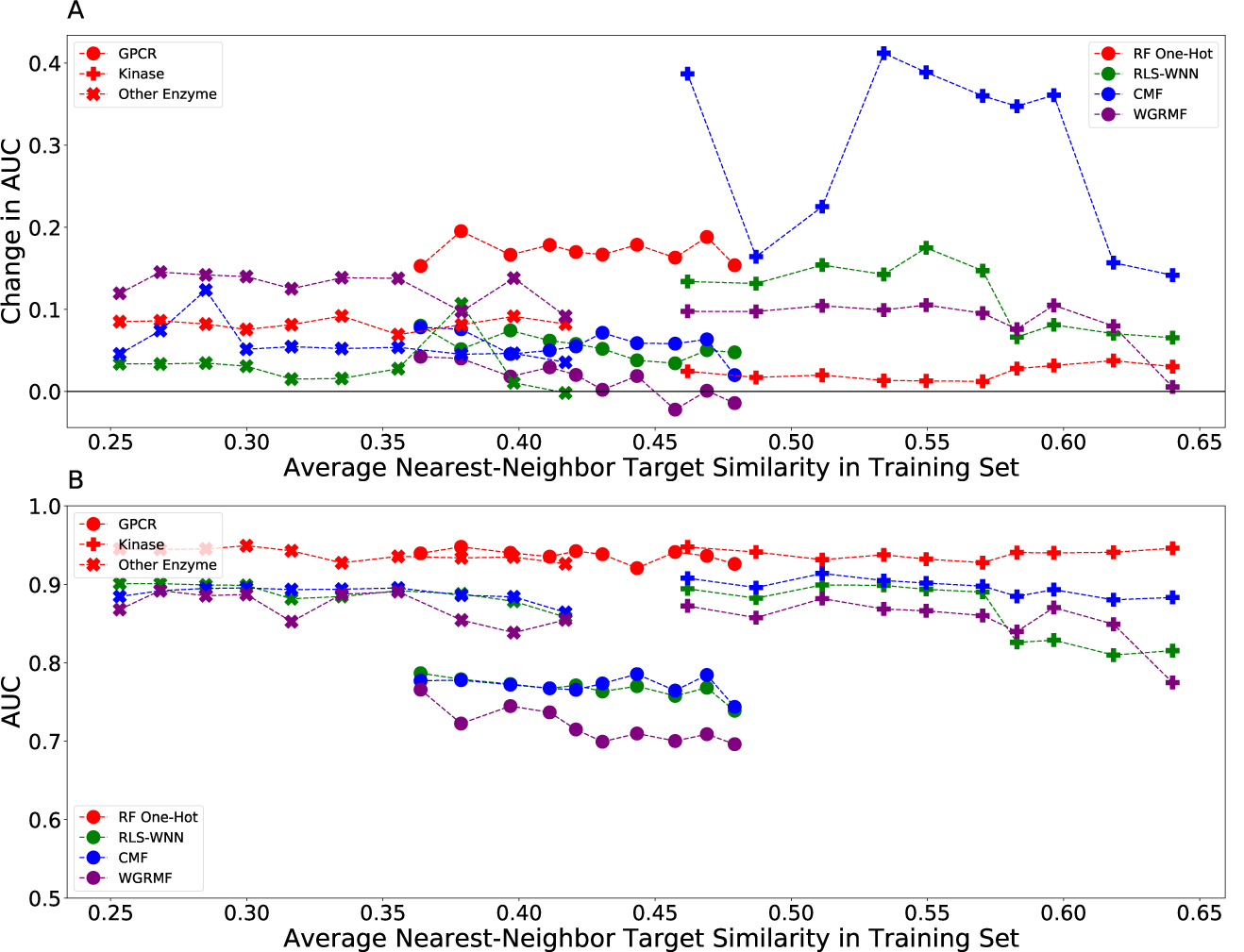
Performance as a Function of Diversity of Training Set, measured by (A) Change in Unseen Drug and Target AUC and (B) Unseen Drug and Target AUC. We measure the diversity of the training set by the average nearest-neighbor similarity between protein targets. Data is shown from 3 datasets: the points are from GPCR, the pluses are from the Kinase dataset, and the dots are from the Ion Channel dataset. As this similarity increases and diversity decreases, our method slightly decreases in effectiveness but continues to improve the unseen target and drug AUC.

## Discussion

In this work, we demonstrate that augmenting the training data for DTI models using predictions made by robust single protein/ligand binding models allows generalisation simultaneously in protein and ligand space so that models can accurately predict interactions between unseen drugs and unseen targets. We observe this effect consistently regardless of dataset size across multiple DTI algorithms. Our best-performing DTI method was a random forest model that uses ECFP6 ligand fingerprints and simple amino acid counts to capture similarity between protein targets. We find that the additional high-quality predictions provided by the single protein binding models significantly improve the accuracy of DTI model predictions. Since this is a very simple model, more complicated models such as deep neural networks will likely also benefit from augmented training data and improve in performance.

Our results suggest that simple feature-based methods such as RF One-Hot perform better than DTI methods that explicitly model the similarities among ligands and protein targets such as CMF. We note that these similarity-based methods implicitly use Tanimoto similarity as the only ligand feature. Analysis of single protein/ligand binding models has indicated that Tanimoto similarity is not a particularly robust metric and options such as logistic regression and random forest learn more (12). Beyond this, the key advantage provided by the data augmentation is simply additional training data. While most interactions are predicted to be inactive, a small but significant proportion of the interactions are predicted to be active (see Figure S12); in contrast, standard approaches either ignore this data or assume that any unknown interactions are inactive (19). We also find that logistic regression consistently performs better for data augmentation than random forest. Figure S12 suggests that this is at least partly a calibration issue; logistic regression outputs probabilities whereas random forests are not generally calibrated (49).

This work raises a number of questions. The DTI models we have used in this paper accept probabilistic training data, but none of them interpret classification probabilities in the most natural way. Cross-entropy loss assumes the output variable is Bernoulli distributed, as appropriate for a classification problem, but all the DTI models use regression approaches and least-squared loss, which assumes the output variable is normally distributed. It is possible that changing the loss function will improve overall performance. Similarly, it is possible that calibrating the single protein/ligand RF models will improve their performance for data augmentation. Furthermore, we note that many of the target similarity-based methods report very similar performance to the baseline methods on the unseen target and drug problem once the single protein/ligand binding models are added. This suggests that the key to improving performance is to develop a better similarity matrix or use explicit features instead, rather than developing more complicated similarity-based models. Finally, an important issue concerns the generalisation ability of both the single target models used to generalise in ligand space and augment the training data, and the DTI models used to generalise in protein space. It is known that single protein/ligand binding models that perform well on benchmark datasets can fail to generalise due to clustering in chemical space and other dataset biases (44–46, 50–53). Similarly, clustering in protein space due to phylogenetic relationships results in similar dataset biases (54) that could result in overly optimistic predictions of generalisability. The high variance in performance between different train/test splits of datasets shown in Figures 2 and 3 suggests that these biases are nontrivial. To drive future improvements, we release the benchmark datasets built for this work. This resource can be leveraged for more careful design of both datasets and data splits to account for these biases, which is an important direction for future work that could yield a better evaluation of these and other models.

## ACKNOWLEDGEMENTS

V.S. acknowledges the support of the Winston Churchill Foundation of the USA. L.J.C. gratefully acknowledges a Next Generation fellowship and support from the Simons Foundation. We acknowledge valuable feedback from Steven Kearnes on an initial draft of this manuscript.

## AUTHOR CONTRIBUTIONS

V.S. and L.J.C.: Development of theoretical framework; statistical modelling; data processing; data analysis. The authors declare no competing interests.

## Methods and Data Availability

### Dataset Construction and Problem Set-up

The datasets presented in this manuscript are built using the same design and methodology as the gold-standard datasets developed by Yamanishi et al. (36). This updated version reflects the vast quantity of protein/ligand binding data that is now available. For these protein targets we found active ligands from ChEMBL 24.1 (55, 56) by filtering for compounds with an IC50, K_*i*_, K_*d*_, or EC50 of less than 1 µM. To prevent duplication we only used proteins from *Homo sapiens*. Targets with fewer than 20 active ligands reported in ChEMBL were eliminated. This resulted in data for 327 protein targets, which we divided into five sub-classes: nuclear receptors, GPCRs, kinases, other enzymes, and ion channels.

Inactive ligands were acquired from two sources: inactive ligands labeled on PubChem indexed by UniProt Protein ID (57, 58) and for targets with a DUD-E decoy set, some inactives from the DUD-E set were included (59). To ensure a reasonable balance of actives to inactives, we also added a randomly selected set of 500 decoys per run; these decoy ligands were selected to not be in any previous set of ligands, either active or inactive. We assumed that these decoys did not interact with any of the given targets. Unlike previous work (19), we did not assume in either training or testing our models that any other unknown interactions between proteins and non-decoy ligands were inactive, since actives for a given protein target may be active for related proteins. Some basic statistics about the size of the datasets may be found in Table 1 and Table S1-3.

**Table 1.**
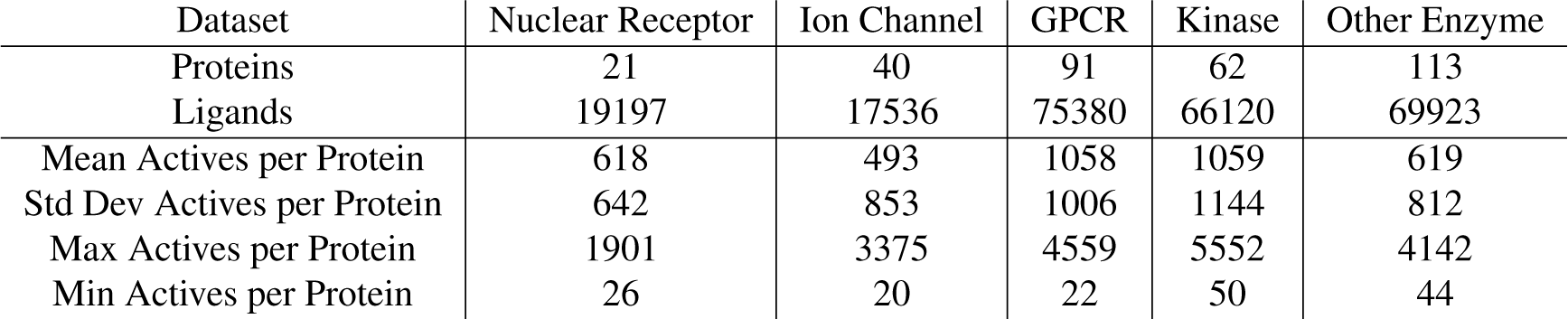
Statistics of Datasets Used. This table reports the number of proteins, ligands, and the mean, standard deviation, maximum, and minimum number of actives per protein. Further information about the datasets can be found in the SI.

We next split each of our five datasets into training and test sets. To explicitly test for generalisation in both protein and ligand space, we used the incomplete training submatrix of protein/ligand interactions as illustrated in Figure 1a. Specifically, we first split the protein targets randomly into an 80% training and 20% test set. For each known target, we randomly selected 20% of the active and known inactive ligands to be in the test set. We further selected 20% of the decoys to be in the test set. All remaining ligands were placed in the training set. The models were not provided with any information about interactions involving any of the test ligands or proteins. Our methodology ensures that all protein targets in the training set have some active and inactive ligands in the training set. Further, the training submatrix is potentially incomplete, since there may be interactions between known proteins and known ligands about which we have no experimental data. A number of statistics about our train/test splits are reported in Table S3.

Our set-up allows us to explicitly test for generalisation in both protein and ligand space by looking at performance on the unseen target and drug subproblem, which tests interactions between test ligands and test proteins. There are 2 other subproblems we may look at: the unseen target subproblem, which looks at interactions between train ligands and test proteins, and the unseen drug subproblem, which looks at interactions between test ligands and train proteins. Our data from Table S1 indicate that despite large variance, on average all 3 subproblems are roughly balanced in terms of the number of actives and inactives due to our setup. This allows us to use the Area Under the Receiver Operating Characteristic Curve (AUC) as our metric for measuring model accuracy.

Some models required hyperparameter tuning, so we created a validation submatrix with the training submatrix using the same methodology as for the train/test split. Hyperparameters were tuned separately for every repetition using the overall AUC on the validation set; model performance was then evaluated on the test set. We performed 20 repetitions to test all models; error bars reported below are to 1 SEM.

### Models

We use logistic regression and random forest with 2048 bit ECFP6 fingerprints generated by rdkit (60) implemented using scikit-learn to build the initial target-specific single protein/ligand binding models. (61, 62) We use regularization constant *C* = 1 for logistic regression and 100 trees with a maximum depth of 25 for random forest; both are known to perform well on the single protein/ligand binding problem.

There are two important modifications that must be made to all DTI models used. First, the DTI model must accept probabilities in its training set. Second, the DTI model must not incorporate any additional information about ligand similarity, since we do not need to generalise to any further ligands. SI Methods Section 2C and Figure S10 show that failing to make this modification results in poorer performance. We made these modifications on a case-by-case basis to all of the models.

We tested weighted nearest neighbor (w-NN) as a baseline DTI model, and four high-performing DTI models from the literature, namely random forest with one-hot features (RF one-hot), regularized least squares (RLS-WNN), collaborative matrix factorization (CMF), and weighted graph-regularized matrix factorization (WGRMF). (19) In all cases, we used pairwise sequence bit scores from pHMMER to describe similarity between protein targets (48), normalized to the length of the amino acid sequence to ensure that each protein’s similarity with itself was 1, and Tanimoto similarities calculated using ECFP6 fingerprints for the ligands. The definitions and properties of the DTI models, and the hyperparameters used for training may be found in SI Methods Sections 1B - 1C.

## Data Availability

Data and code required to reproduce the results of this paper are available in the Supporting Information.

