## Supplementary material for "Using Single Protein/Ligand Binding Models to Predict Active Ligands for Unseen Proteins": SI Appendix

#### This PDF file includes:

Supplementary text
Figs. S1 to S16
Tables S1 to S3
References for SI reference citations

### Supporting Information Text

### 1. Supplementary Methods

**A. Dataset Statistics.** Some basic statistics that describe the five protein datasets built for this study may be found in Table S1. For details about how these five datasets were built, please see the Methods section of the main text. Scripts that can be used to rebuild and update these datasets are provided as part of the Supplementary Code provided with this study. Statistics that characterize the diversity of the protein targets, ligands and active ligands for each dataset are provided in Table S2. Here the similarity across the protein targets in each dataset is calculated as described in the Methods section of the main text. Statistics for ligand similarity are calculated using Tanimoto similarities and ECFP6 fingerprints across (i) all ligands from each dataset, and (ii) all active ligands from each dataset. Finally, in Table S3 we report the size of the train and test sets used for each dataset, in addition to the mean similarity between any protein (or ligand) in the test set and the closest protein (or ligand) in the corresponding training set.

**Table S1. Data Statistics. We report the number of proteins, ligands, active interactions and non-decoy inactive interactions for each of the five protein target datasets constructed in this study together with statistics describing the active ligands per target.**

| Dataset | Nuclear Receptor | Ion Channel | GPCR | Kinase | Other Enzyme |
|---|---|---|---|---|---|
| Proteins | 21 | 40 | 91 | 62 | 113 |
| Ligands | 19197 | 17536 | 75380 | 66120 | 69923 |
| Actives | 12978 | 19717 | 96289 | 65654 | 69968 |
| Non-Decoy Inactives | 11438 | 1480 | 13289 | 28899 | 16972 |
| Mean Actives per Protein | 618 | 493 | 1058 | 1059 | 619 |
| Std Dev Actives per Protein | 642 | 853 | 1006 | 1144 | 812 |
| Max Actives per Protein | 1901 | 3375 | 4559 | 5552 | 4142 |
| Min Actives per Protein | 26 | 20 | 22 | 50 | 44 |

**Table S2. Dataset Similarity Statistics. We calculate statistics describing the similarity between all proteins in each dataset, between all ligands in each dataset, and between all ligands labelled active for the same target. Here NN refers to nearest-neighbor similarity, or similarity to the nearest neighbor of a given point.**

| Dataset | Nuclear Receptor | Ion Channel | GPCR | Kinase | Other Enzyme |
|---|---|---|---|---|---|
| Mean Protein Similarity | 0.145 | 0.033 | 0.074 | 0.081 | 0.009 |
| Std Dev Protein Similarity | 0.125 | 0.121 | 0.074 | 0.104 | 0.056 |
| Mean NN Protein Similarity | 0.48 | 0.42 | 0.38 | 0.48 | 0.34 |
| Std NN Protein Similarity | 0.21 | 0.28 | 0.15 | 0.23 | 0.28 |
| Mean Ligand Similarity | 0.084 | 0.093 | 0.094 | 0.092 | 0.088 |
| Std Dev Ligand Similarity | 0.036 | 0.042 | 0.032 | 0.032 | 0.032 |
| Mean NN Ligand Similarity | 0.61 | 0.71 | 0.72 | 0.66 | 0.67 |
| Std Dev NN Ligand Similarity | 0.20 | 0.15 | 0.15 | 0.18 | 0.17 |
| Mean Active Ligand Similarity | 0.122 | 0.125 | 0.133 | 0.138 | 0.136 |
| Std Dev Active Ligand Similarity | 0.085 | 0.089 | 0.082 | 0.098 | 0.087 |
| Mean NN Active Ligand Similarity | 0.70 | 0.72 | 0.74 | 0.72 | 0.69 |
| Std Dev NN Active Ligand Similarity | 0.17 | 0.14 | 0.13 | 0.16 | 0.17 |

**Table S3. Data Splits. We report the number of training and test ligands, and the mean similarity between any protein or ligand in the test set and the closest protein or ligand in the corresponding training set. We also report the number of actives and inactives in the unseen drug, unseen target, and unseen drug + target problems. All numbers listed are mean $\pm$ standard deviation, where the mean is taken across $20$ different train-test splits.**

| Dataset | Nuclear Receptor | Ion Channel | GPCR | Kinase | Other Enzyme |
|---|---|---|---|---|---|
| Train Ligands | $14969 \pm 14$ | $13860 \pm 14$ | $55885 \pm 32$ | $49776 \pm 45$ | $53865 \pm 28$ |
| Test Ligands | $4723 \pm 14$ | $4172 \pm 14$ | $19974 \pm 32$ | $16830 \pm 45$ | $16537 \pm 28$ |
| Mean Train-Test Protein Similarity | $0.51 \pm 0.20$ | $0.43 \pm 0.28$ | $0.37 \pm 0.14$ | $0.47 \pm 0.23$ | $0.33 \pm 0.28$ |
| Mean Train-Test Ligand Similarity | $0.61 \pm 0.20$ | $0.70 \pm 0.16$ | $0.72 \pm 0.15$ | $0.67 \pm 0.18$ | $0.67 \pm 0.17$ |
| Unseen Drug Actives | $3018 \pm 350$ | $4168 \pm 561$ | $25979 \pm 1170$ | $20492 \pm 1125$ | $16083 \pm 1374$ |
| Unseen Drug Inactives | $3861 \pm 290$ | $3508 \pm 54$ | $10250 \pm 375$ | $10860 \pm 688$ | $12306 \pm 315$ |
| Unseen Target Actives | $2020 \pm 932$ | $2927 \pm 1426$ | $13658 \pm 2252$ | $8417 \pm 2455$ | $10098 \pm 2731$ |
| Unseen Target Inactives | $4185 \pm 841$ | $3407 \pm 165$ | $8957 \pm 1548$ | $9561 \pm 2261$ | $11503 \pm 1099$ |
| Unseen Target + Drug Actives | $889 \pm 351$ | $1116 \pm 567$ | $7127 \pm 1115$ | $5435 \pm 1071$ | $4384 \pm 1379$ |
| Unseen Target + Drug Inactives | $1275 \pm 285$ | $889 \pm 49$ | $2655 \pm 378$ | $2866 \pm 696$ | $3207 \pm 301$ |

**B. Model Descriptions.** A list of the DTI models tested follows, along with the modifications we made. For convenience, let the training drug-target similarity matrix be $Y$, the predicted output from the model $Y_{\text{pred}}$, the drug similarity matrix be $S_d$, and the target similarity matrix be $S_t$.

1. Weighted Nearest Neighbor (Weighted NN), a baseline method which weights the nearest interactions in protein and ligand space by their similarity (1). Unknown interactions are assumed to be non-interactions.

2. Random Forest One-Hot (RF One-Hot). The feature set used was ECFP6 fingerprints with 2048 bits for the ligands and a one-hot count of amino acids for the proteins. This is distinct from the single protein/ligand binding models since it attempts to fit the data for all the proteins at once. In order to incorporate the probabilistic information, we used a regressor model instead of a classifier with a least-squared loss.

3. Regularized Least Squares-Weighted Nearest Neighbor (RLS-WNN), a regularized least-squares linear regression model based on protein and ligand similarity. Unknown interactions are assumed to be non-interactions. We first infer interactions for completely unseen drugs and targets by letting

$$Y(d_i) = \sum_{j=1}^{n} w_j Y(d_j)$$

   where the drugs are sorted in descending order of similarity and $w_j = \eta^{j-1}$ for some hyperparameter $\eta$ (2).

   Next, we compute Gaussian interaction profile (GIP) kernels; the drug kernel is

$$\text{GIP}_d(d_i, d_j) = \exp(-\gamma ||Y(d_i) - Y(d_j)||^2)$$

   for some hyperparameter $\gamma$ and the target kernel is defined similarly. We then merge these with the similarity matrices to get

$$K_d = \alpha S_d + (1 - \alpha)\,\text{GIP}_d$$

   for some hyperparameter $\alpha, 0 \leq \alpha \leq 1$. Let $K = K_d \otimes K_t$ be the kernel over drug-target pairs; we can then use kernel ridge regression to compute

$$\hat{Y}_{\text{pred}} = K(K + \sigma I)^{-1} \hat{Y}$$

   where $\hat{Y}$ is the flattened version of the training matrix. The inverse is computed via an eigendecomposition due to computational constraints (2).

   Due to the larger dataset size, we approximated the inverse of the drug similarity matrix using only the top 100 eigenvalues instead of computing it exactly. When we incorporated our data augmentation step, we only used the target similarity matrix in the RLS computation and omitted the weighted nearest neighbor preprocessing.

4. Collaborative Matrix Factorization (CMF) based on protein and ligand similarity. CMF finds matrices $A$ and $B$ to minimize the function

$$\left|\left|W \odot \left(Y - AB^T\right)\right|\right|^2 + \lambda_l \left(||A||^2 + ||B||^2\right) + \lambda_t \left|\left|S_t - AA^T\right|\right|^2 + \lambda_d \left|\left|S_d - BB^T\right|\right|^2$$

   where $W$ is a weight matrix with $W_{ij} = 0$ for unknown pairs, $\odot$ is an elementwise product, all norms are the Frobenius norm, and $\lambda_l$, $\lambda_d$, and $\lambda_t$ are hyperparameters. Optimization occurs via an alternating least squares algorithm (3). When incorporating the single protein/ligand binding models, we set the hyperparameter $\lambda_d$ corresponding to drug similarity to 0.

5. Weighted Graph-Regularized Matrix Factorization (WGRMF) based on protein and ligand similarity. The loss function is

$$\left|\left|W \odot \left(Y - AB^T\right)\right|\right|^2 + \lambda_l \left(||A|^2 + ||B||^2\right) + \lambda_t \,\text{Tr}\left(A^T \ell_t A\right) + \lambda_d \,\text{Tr}\left(B^T \ell_d B\right)$$

   where $\ell_d, \ell_t$ are normalized graph Laplacians for a sparsified drug/target graph. Sparsification occurred by only keeping the $p$ nearest neighbors for any given point in the graph. Optimization is essentially the same as in CMF (4). When incorporating the single protein/ligand binding models, we set the hyperparameter corresponding to drug similarity to 0.

**C. Hyperparameter Selection.** All hyperparameters were individually selected for each run of every given model. In order to select the hyperparameters, we used a 3-way train/validation/test split, where the train+validation submatrix was generated as described in the main text and the training submatrix was generated as a subset of the train+validation submatrix. Specifically, we used the procedure previously described to identify train+validation and test ligands and proteins. We then repeated this procedure within the set of train+validation ligands and proteins, splitting the protein targets randomly into an 75% training and 25% test set and proceeding similarly with the ligands. The optimal hyperparameter on the validation set was selected and used to measure performance on the test set.

Hyperparameters were adapted from successful values in the literature (5). The list of hyperparameters used follows:

1. Weighted NN had no relevant hyperparameters.

2. RF One-Hot. The number of trees was allowed to vary within $\{50, 100, 150\}$ and the maximum depth $\{15, 20, 25, 30\}$.

3. RLS-WNN. The ordered triple $(\sigma, \alpha, \eta)$ was allowed to vary within

$$\{(0.25, 0.1, 0.4), (0.5, 0.9, 0.4), (2, 0.6, 0.1), (0.5, 1, 0.6)\}.$$

When preprocessing was added, the same ordered triple was allowed to vary within

$$\{(0.5, 0.5, 0.6), (0.5, 1, 0.7), (0.25, 1, 0.7)\};$$

we found these values consistently performed better than the previous ones only with preprocessing.

4. CMF. The ordered triple $(\lambda_l, \lambda_d, \lambda_t)$ was allowed to vary within

$$\{(1, 4, 32), (2, 8, 0.125), (4, 32, 0.25), (2, 64, 0.125), (0.25, 0.25, 32), (1, 0.0625, 2)\}.$$

When preprocessing was added, we fixed $\lambda_d = 0$ and allowed

$$(\lambda_l, \lambda_t) \in \{(1, 32), (0.25, 32), (0.5, 64), (0.5, 32)\}.$$

5. WGRMF. The ordered quadruple $(\lambda_l, \lambda_d, \lambda_t, p)$ was allowed to vary within

$$\{(0.0625, 0.05, 0.1, 2), (0.25, 0.2, 0.2, 4), (0.25, 0.2, 0.2, 4), (0.25, 0.2, 0.2, 5)\}.$$

When preprocessing was added, we fixed $\lambda_d = 0$ in the above values. $p$ determines the degree of sparsification of the training matrix used as a form of regularization prior to running the WGRMF algorithm (4).

### 2. Supplementary Results

**A. Additional Models and Subproblems.** Using our setup, we can define 3 different subproblems: the unseen target and drug subproblem consists of test drugs and targets both without experimental data, the unseen target subproblem consists of train drugs with experimental data and test targets without, and the unseen drug subproblem consists of train targets with experimental data and test drugs without. The main text focused on the unseen target and drug subproblem; we now examine model performance on all 3.

Figures S2, S3, and S4 show the improvement in performance on the unseen drug, the unseen target, and the both unseen drug and target subproblems upon adding single protein/ligand binding models for all DTI models, not just the 3 shown in the main text. We see consistent improvement statistically significant at a $p < 0.01$ level regardless of the size or other properties of the dataset and improvement for both logistic regression and random forest. The only exception is RF one-hot on the unseen target problem; it is unclear why this is. We see improvement even for models such as WGRMF, where adding single protein/ligand binding models has minimal effect on the unseen drug subproblem but a much larger effect in the unseen target subproblem. Thus even a model that performs well in the unseen drug subproblem can benefit from our data augmentation method.

**B. Trial-by-Trial Comparisons.** Instead of examining averages over all 20 trials of the algorithm, we can also examine performance on each individual train/test split. A comparison of the unseen drug and target AUCs is shown in Figure S5, of the unseen target AUCs in Figure S6, and of the unseen drug AUCs in Figure S7. Each data point in these plots corresponds to one of the 20 train/validation splits and shows the AUC of our models on the appropriate subproblem with and without logistic regression.

Figures S5 and S6 demonstrate that despite large variance in performance between trials, on almost every individual trial for almost all models we see consistent improvement. The lone exception is RF One-Hot on the unseen target AUC, as mentioned above. The variance in model performance is likely due to randomness in the set of test proteins; since there are a small number of protein targets for all datasets, if a protein that is particularly distant is held-out, that could decrease performance. This variance also tends to decrease after application of our method.

Figure S7 demonstrates that our models consistently perform outstandingly on the unseen drug subproblem. As noted in the main text, this is likely at least partially a result of dataset bias in chemical space.

We make the same comparisons with random forest used as the single protein/ligand binding model in Figures S8, S9, and S10. Our conclusions are similar as to the case with logistic regression.

**C. Failure to Eliminate Ligand Similarity.** To better understand the importance of eliminating the ligand similarity matrix in the DTI model, we examined the effects of not eliminating it in the RLS-WNN algorithm. We compared five models: the original RLS-WNN algorithm, the RLS-WNN algorithm unmodified but with logistic regression or random forest, and the RLS-WNN modified to only look at the target similarity matrix.

Our results are shown in Figure S11. We observe that the unmodified RLS-WNN algorithm with a single protein/ligand binding model is often better than the original RLS-WNN model, but it is still significantly worse than the RLS-WNN model that correctly ignores ligand similarity. In particular, the unseen target AUCs and some of the unseen drug AUCs are significantly worse without the modification. We also see that when the unseen drug AUC is not significantly affected like in the nuclear receptor dataset, the drop in performance in the other AUCs is smaller; however, when the unseen drug AUC is significantly affected, the drop in performance in the other AUCs is larger, almost negating the advantages of our method.

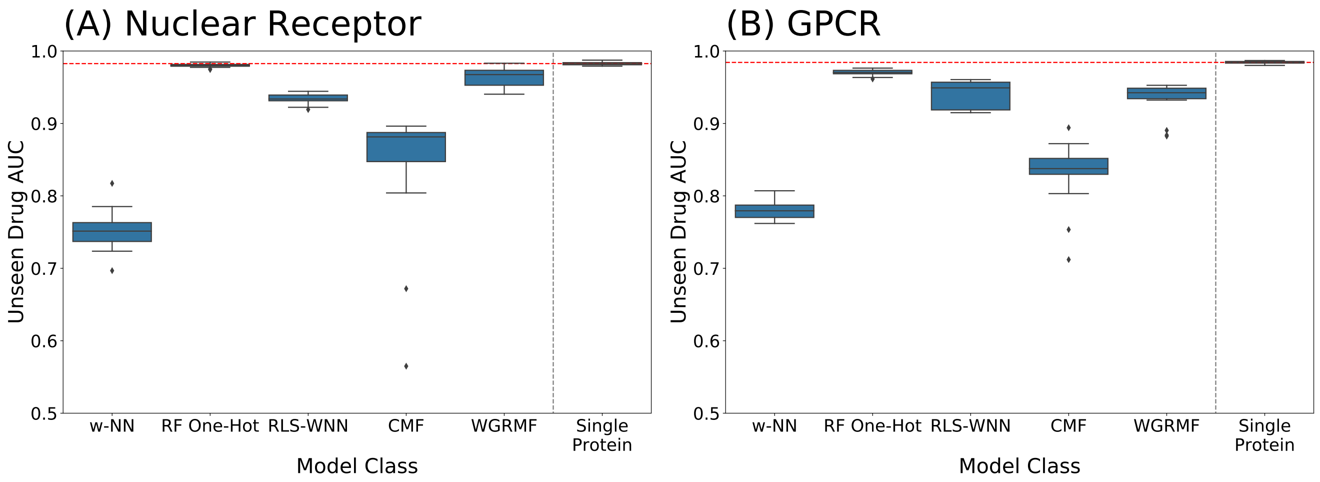


**Fig. S1.** Performance of Single Protein Models and DTI Models for protein targets with available training data from (a) the Nuclear Receptor Dataset and (b) the GPCR Dataset. Despite the additional information about related proteins available to the DTI models, the single protein/ligand binding models consistently outperform them at predicting whether unseen ligands bind to protein targets from the training set. This motivates using the single protein/ligand binding models to improve performance of DTI models at making predictions for the harder case or protein targets for which no training data is available.

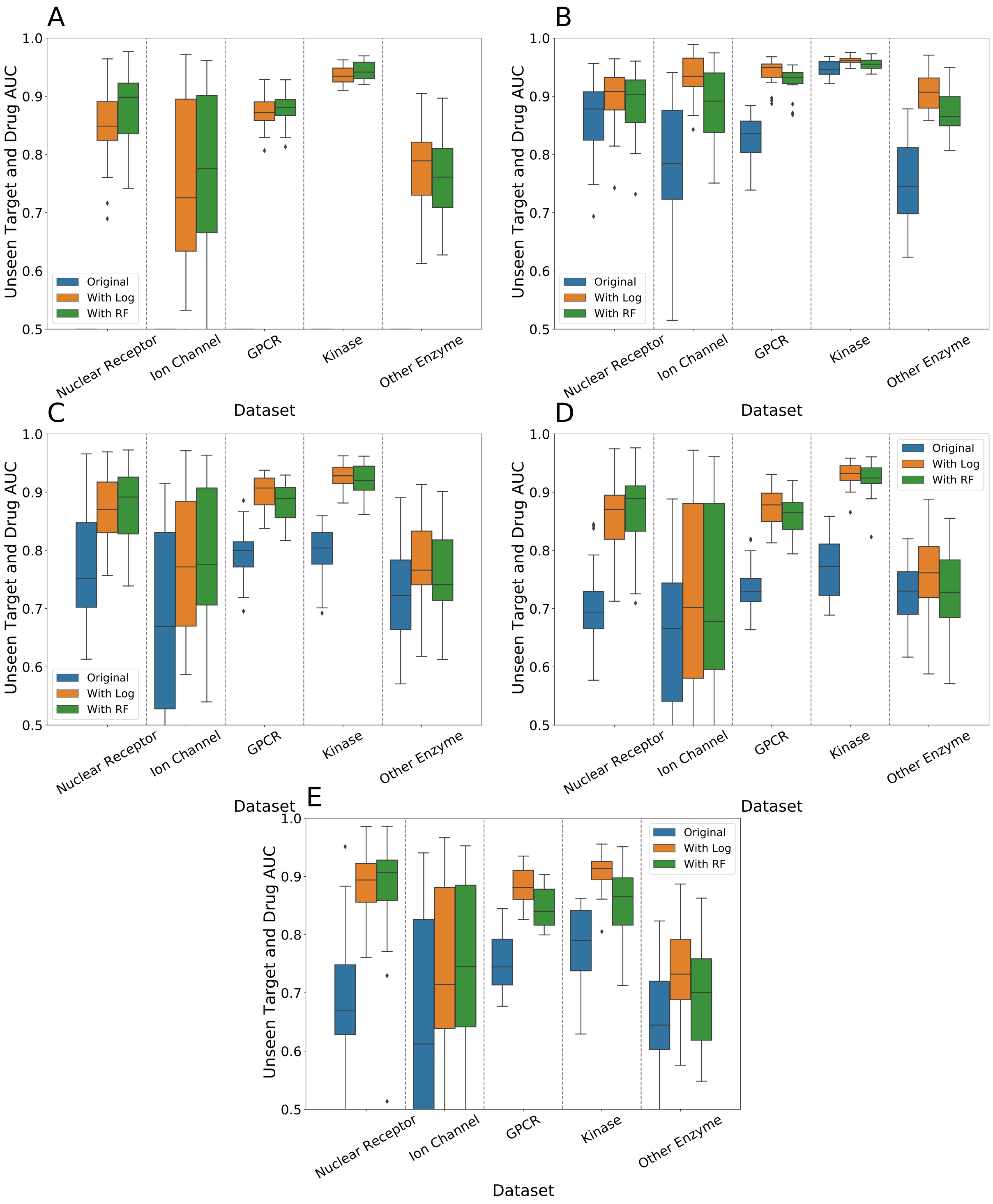


**Fig. S2.** Unseen Target and Drug AUCs for (a) weighted NN, (b) RF one-hot, (c) RLS-WNN, (d) CMF, and (e) WGRMF. We see consistent improvement in all models upon adding the data from single protein/ligand binding models; logistic regression performs slightly better than random forest.

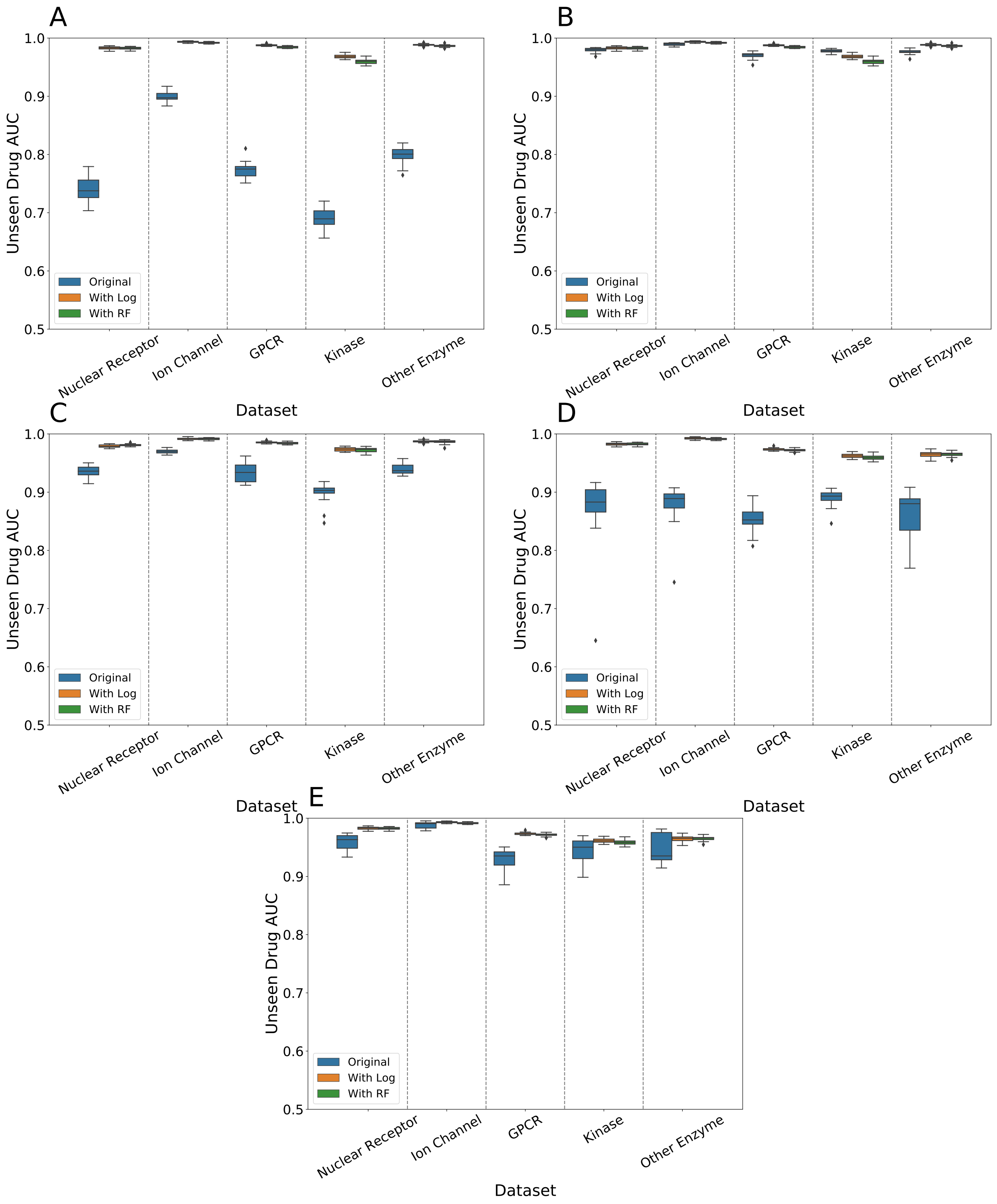


**Fig. S3.** Unseen Drug AUCs for (a) weighted NN, (b) RF one-hot, (c) RLS-WNN, (d) CMF, and (e) WGRMF. We see consistent improvement in all models upon adding the data from single protein/ligand binding models.

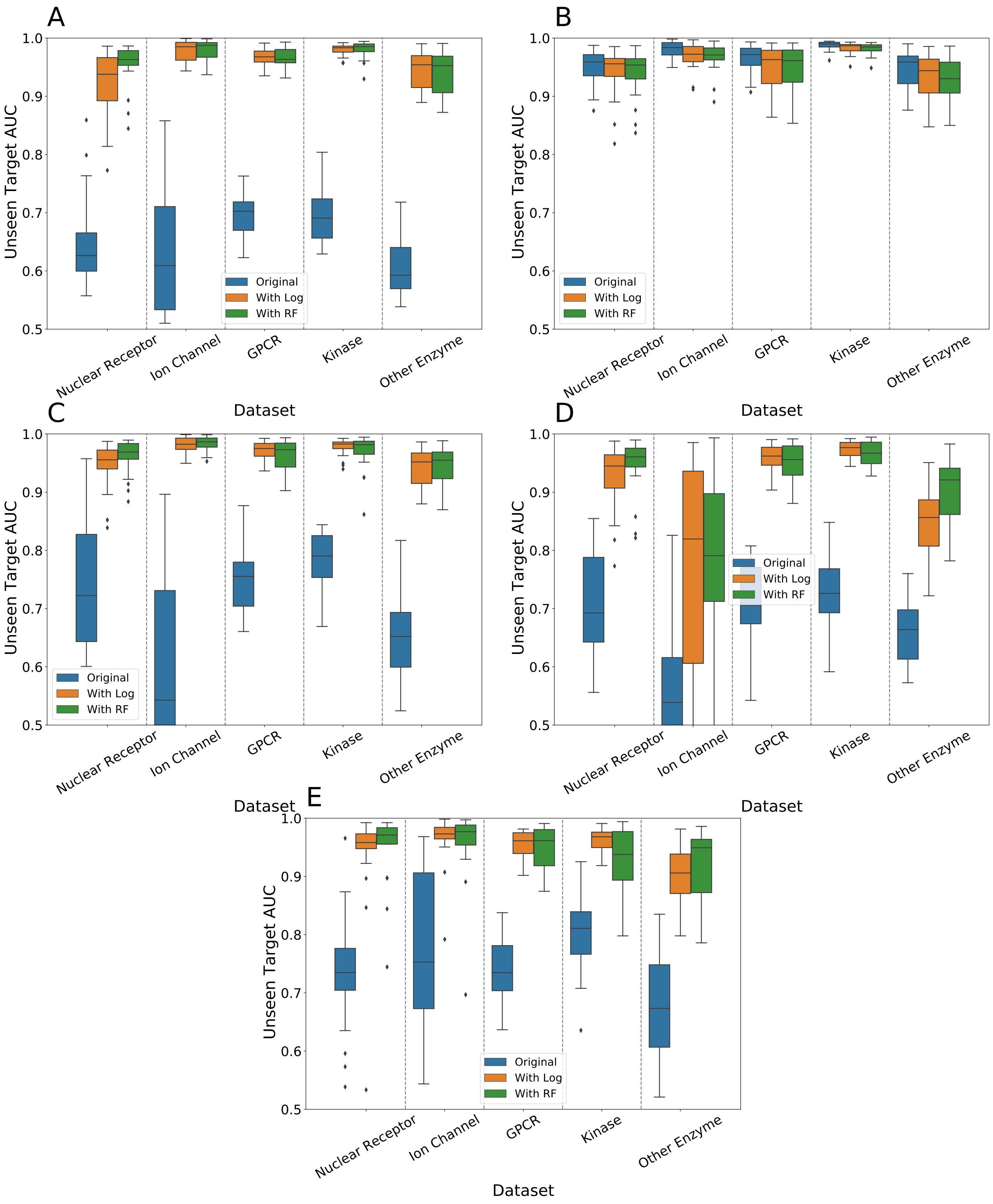


**Fig. S4.** Unseen Target AUCs for (a) weighted NN, (b) RF one-hot, (c) RLS-WNN, (d) CMF, and (e) WGRMF. We see consistent improvement in all models aside from RF one-hot upon adding the data from single protein/ligand binding models.

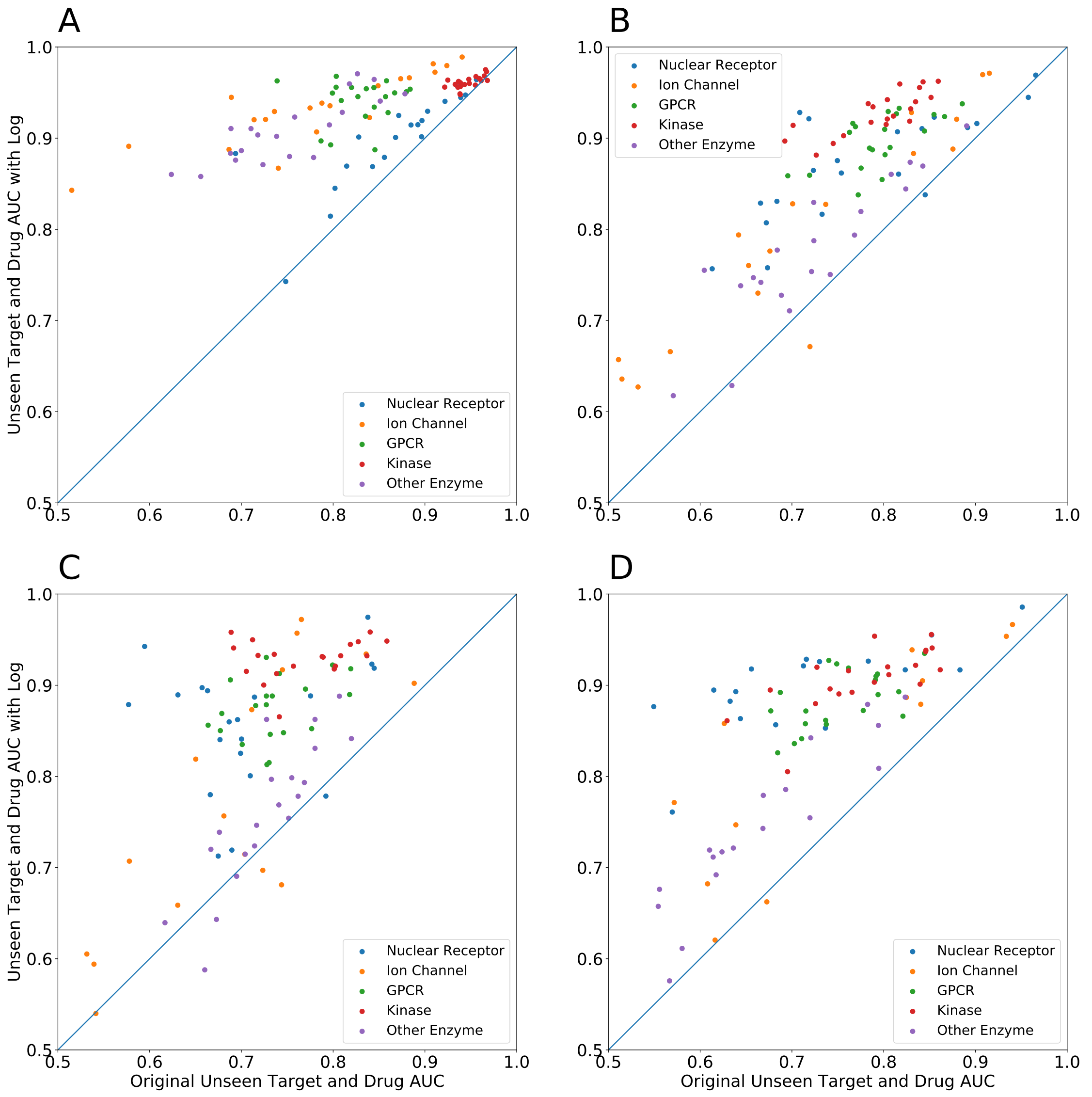


**Fig. S5.** Comparison of Unseen Drug and Target AUCs with and without Logistic Regression for (a) RF One-Hot, (b) RLS-WNN, (c) CMF, and (d) WGRMF. We see improvement on every method in almost all test runs. The wide variance in performance is likely due to randomness in which proteins are held-out; our method also decreases this variance.

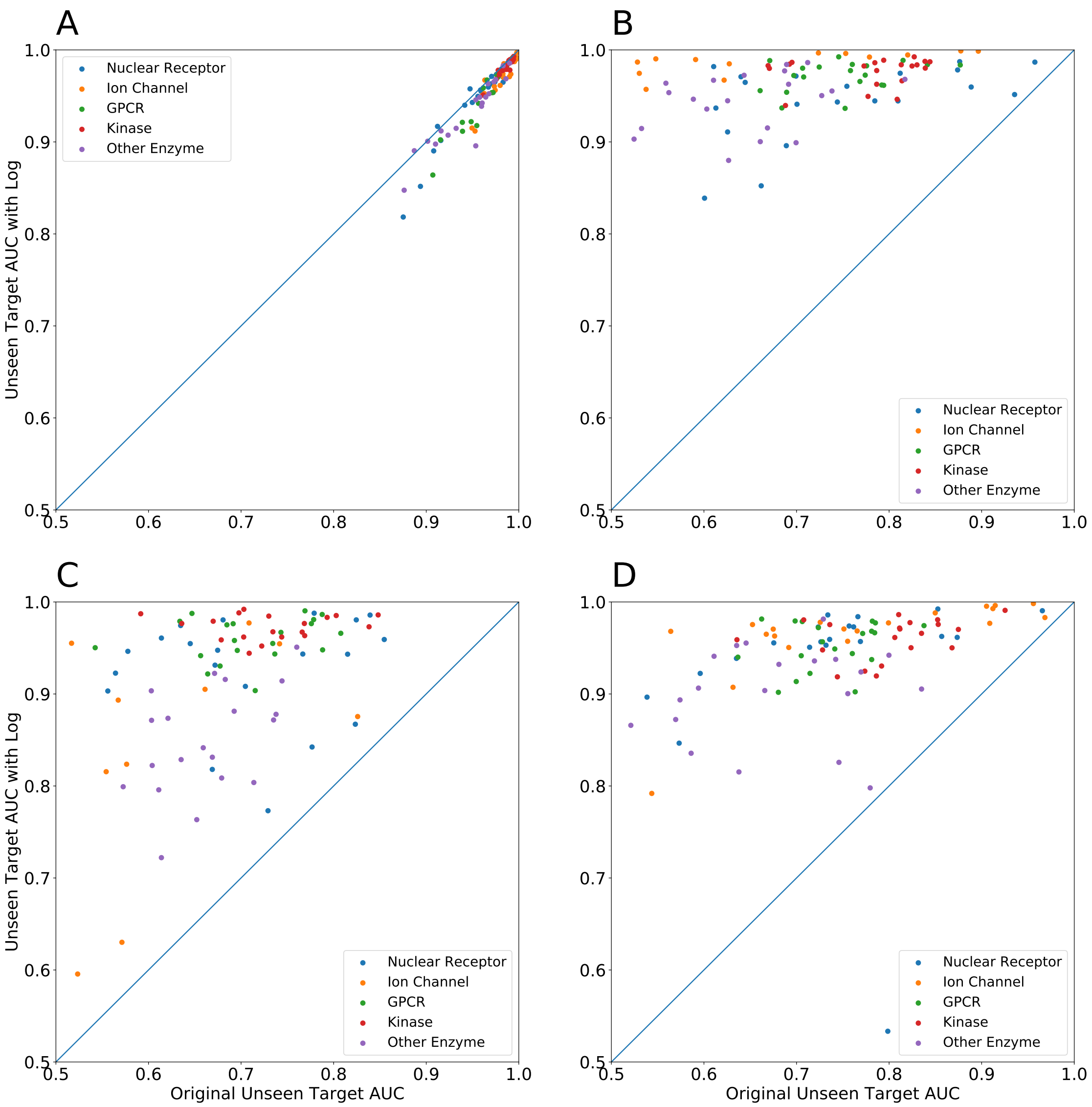


**Fig. S6.** Comparison of Unseen Target AUCs with and without Logistic Regression for (a) RF One-Hot, (b) RLS-WNN, (c) CMF, and (d) WGRMF. We see improvement on every method in almost all test runs for models aside from RF One-Hot. The wide variance in performance is likely due to randomness in which proteins are held-out; our method also decreases this variance.

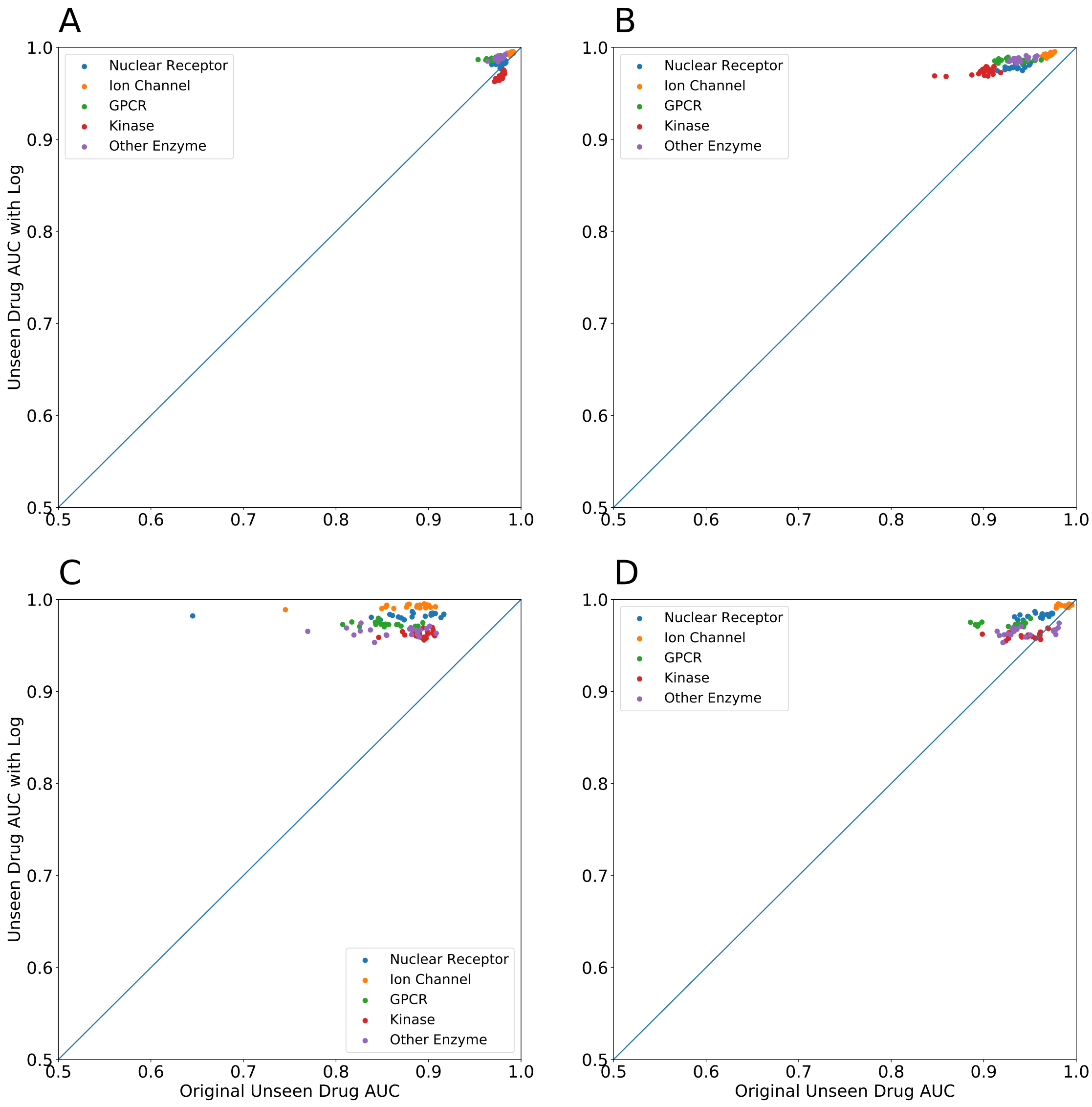


**Fig. S7.** Comparison of Unseen Drug AUCs with and without Logistic Regression for (a) RF One-Hot, (b) RLS-WNN, (c) CMF, and (d) WGRMF. We see improvement on every method in almost all test runs. The high performance is likely a result of bias within the chemical dataset.

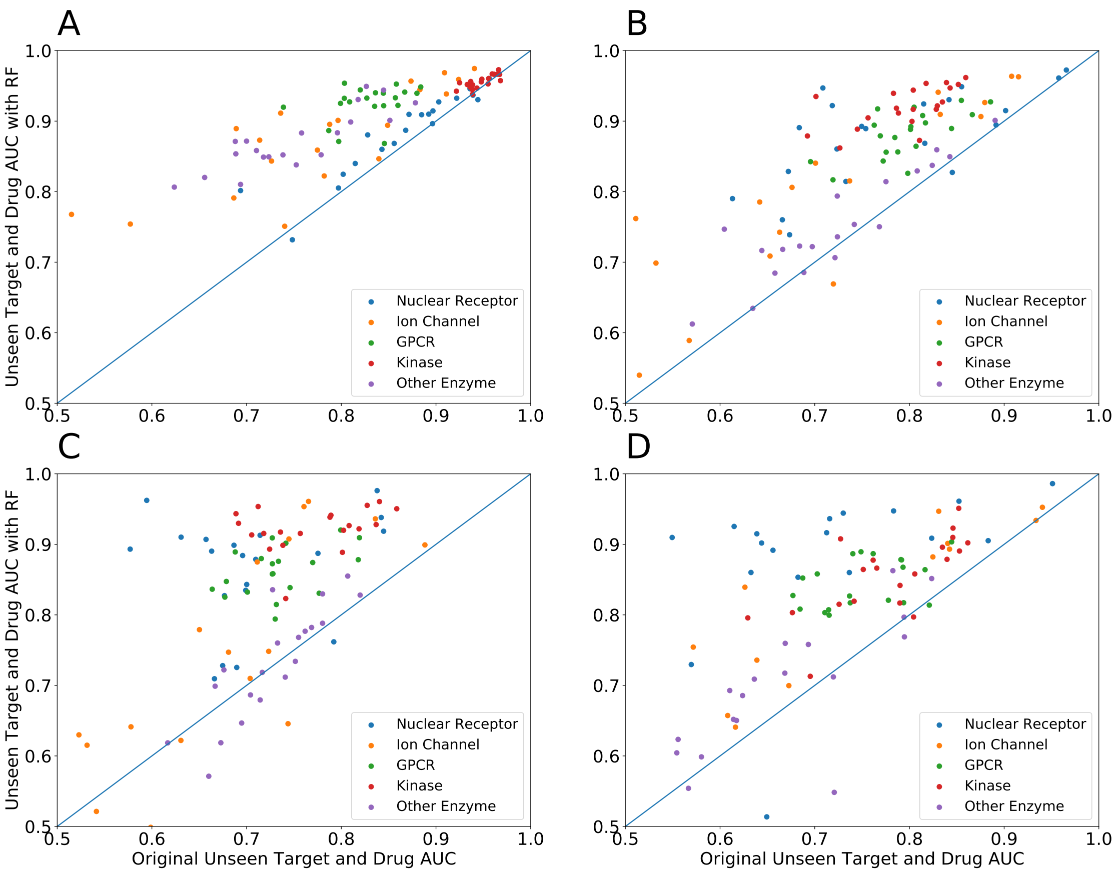


**Fig. S8.** Comparison of Unseen Drug and Target AUCs with and without Random Forest for (a) RF One-Hot, (b) RLS-WNN, (c) CMF, and (d) WGRMF. We see improvement on every method in almost all test runs. The wide variance in performance is likely due to randomness in which proteins are held-out; our method also decreases this variance.

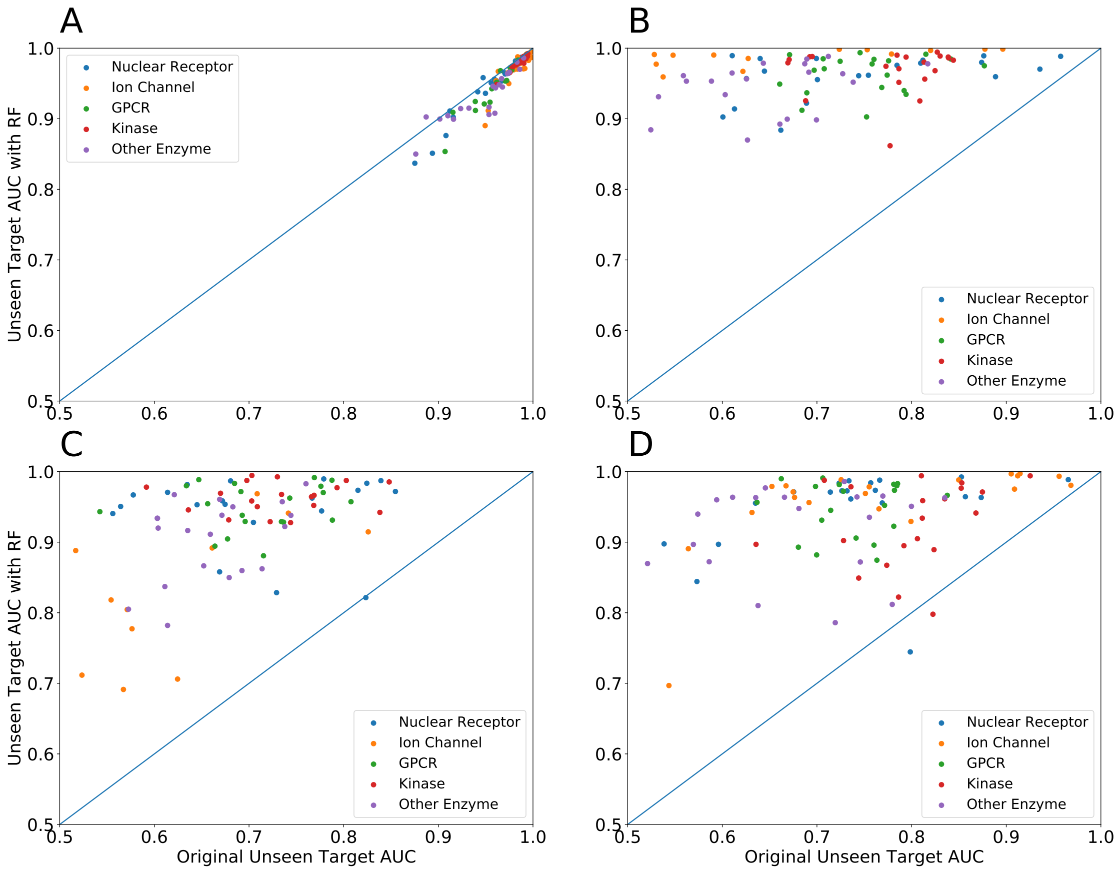


**Fig. S9.** Comparison of Unseen Target AUCs with and without Random Forest for (a) RF One-Hot, (b) RLS-WNN, (c) CMF, and (d) WGRMF. We see improvement on every method in almost all test runs for models aside from RF One-Hot. The wide variance in performance is likely due to randomness in which proteins are held-out; our method also decreases this variance.

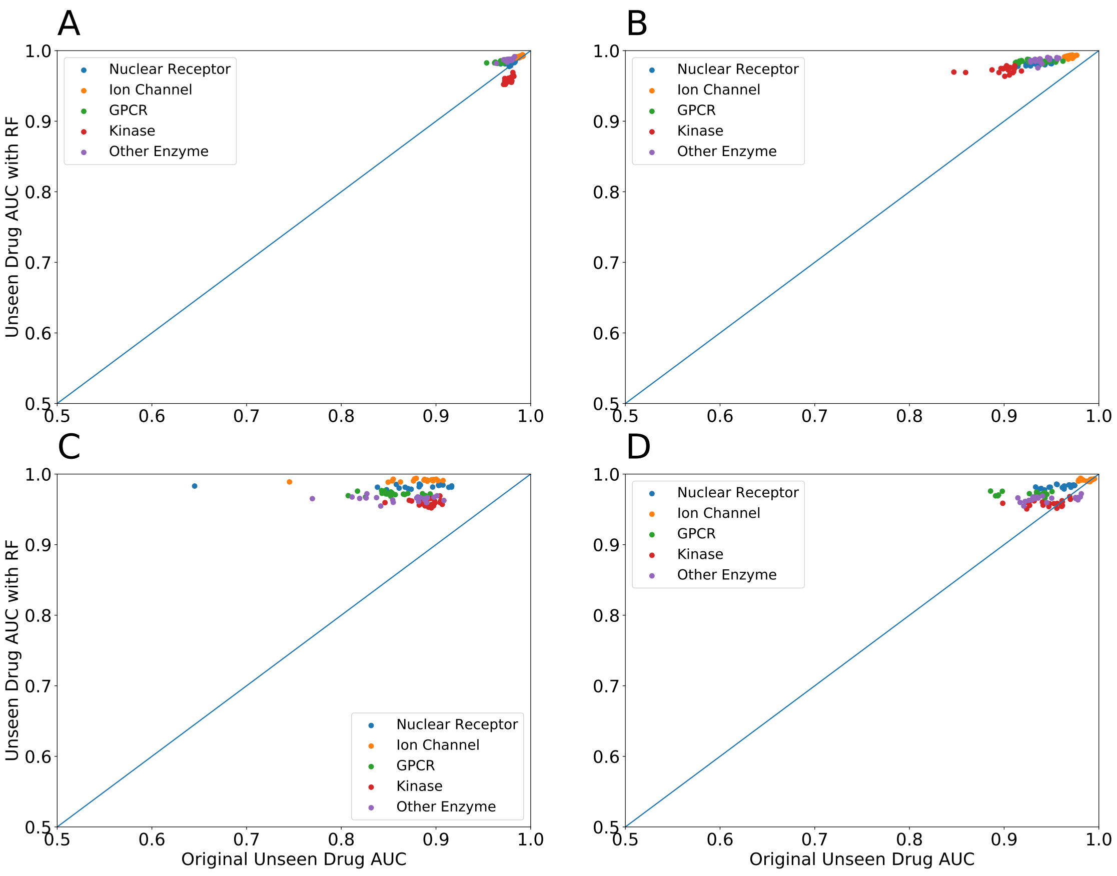


**Fig. S10.** Comparison of Unseen Drug AUCs with and without Random Forest for (a) RF One-Hot, (b) RLS-WNN, (c) CMF, and (d) WGRMF. We see improvement on every method in almost all test runs. The high performance is likely a result of bias within the chemical dataset.

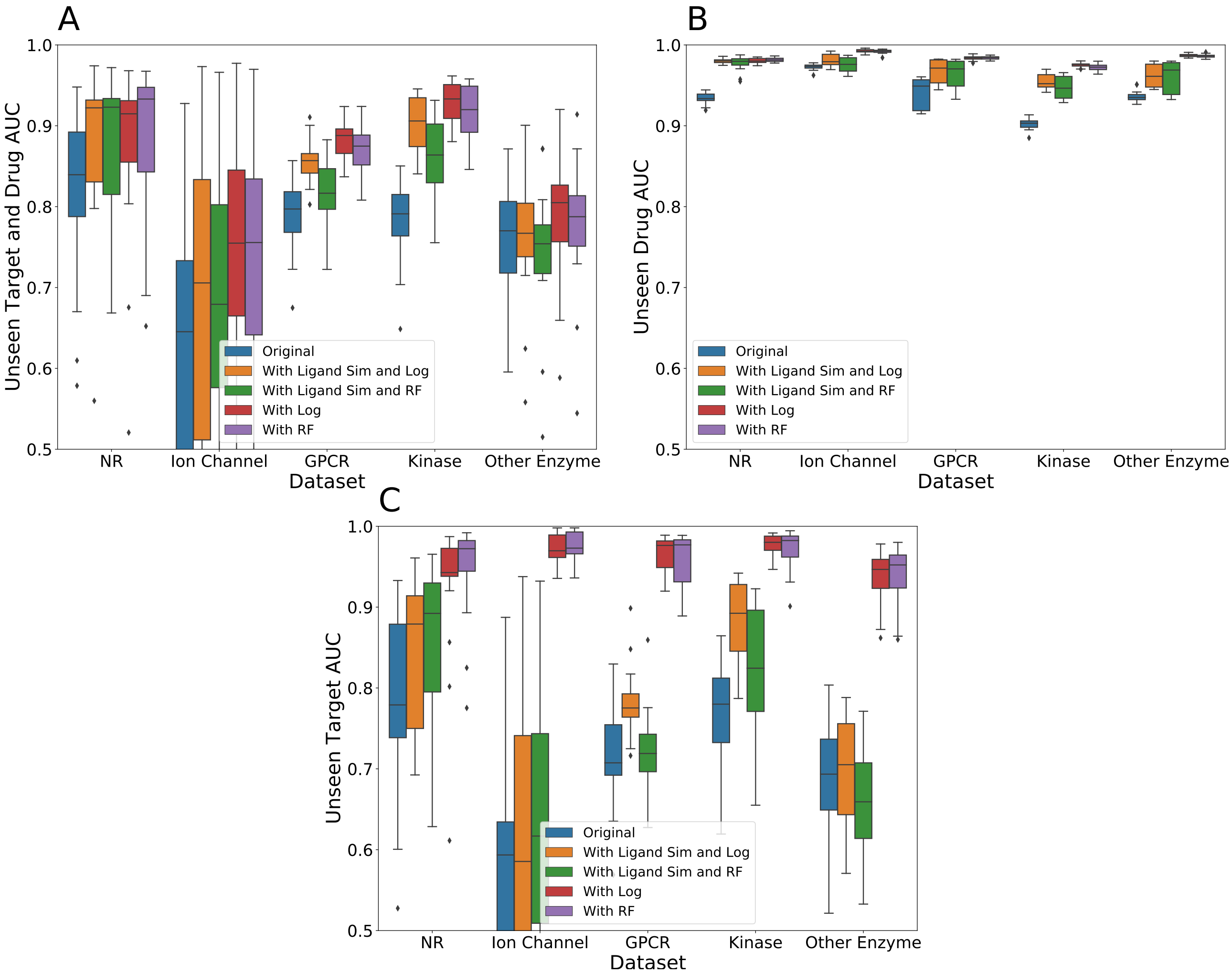


**Fig. S11.** Effect of Not Eliminating Ligand Similarity on RLS for (a) Unseen Target and Drug AUC, (b) Unseen Drug AUC, and (c) Unseen Target AUC. Models that incorrectly incorporate ligand similarity into the RLS algorithm consistently perform worse than those that do not, emphasizing the need to eliminate ligand similarity in the DTI model after using a single protein/ligand binding method.

Clearly, adding information about ligand similarity to the DTI model worsens the performance of the overall model. This is likely because logistic regression and random forest are more accurate methods of determining similarity between ligands than the Tanimoto similarity matrix. Incorporation of the Tanimoto ligand similarity matrix corrupts the information provided by the single protein/ligand binding method and lowers accuracy of the models overall. It is therefore important to ensure that DTI methods do not incorporate additional inaccurate information about the ligands; when our method is used, they should be used exclusively for generalisation in target space.

**D. Probabilities Predicted in Training Submatrix.** As mentioned in the main text, one of the primary explanations behind the added predictive power of our method is the additional information provided by filling in the training matrix for all of the train proteins. We show a histogram of the probabilities of interaction predicted by both logistic regression and random forest in Figure S12 for all datasets.

We see a number of active interactions predicted that were not in the original training matrix. This added information helps the DTI models train more effectively and make more accurate predictions. We also see issues with random forest calibration, suggesting that recalibrating the random forest model could improve overall performance.

**E. Performance Analysis.** We also examined the dependence of model performance on other factors, like the number of proteins in the training set. To do this, we generated train/test splits of the data by the same methodology as used previously, but with varying proportions of the protein targets in the training set. We found in Figure S13 that model performance showed no particularly strong dependence on the size of the training set. As mentioned in the main text, we similarly found no strong dependence on the diversity of the training set.

Figure S14 shows the relationship between the number of known active ligands per training protein and model performance. This data was acquired over the same trials used to generate Figures 2 and 3 in the main text. We see no particularly strong dependence on the number of known active ligands per protein, but we did not experiment with extreme scenarios where very few active ligands are known for a given protein.

To provide further context for Figures 3 and 4, Figures S15 and S16 show the distribution of target and ligand similarity, respectively, across all 20 trials run.

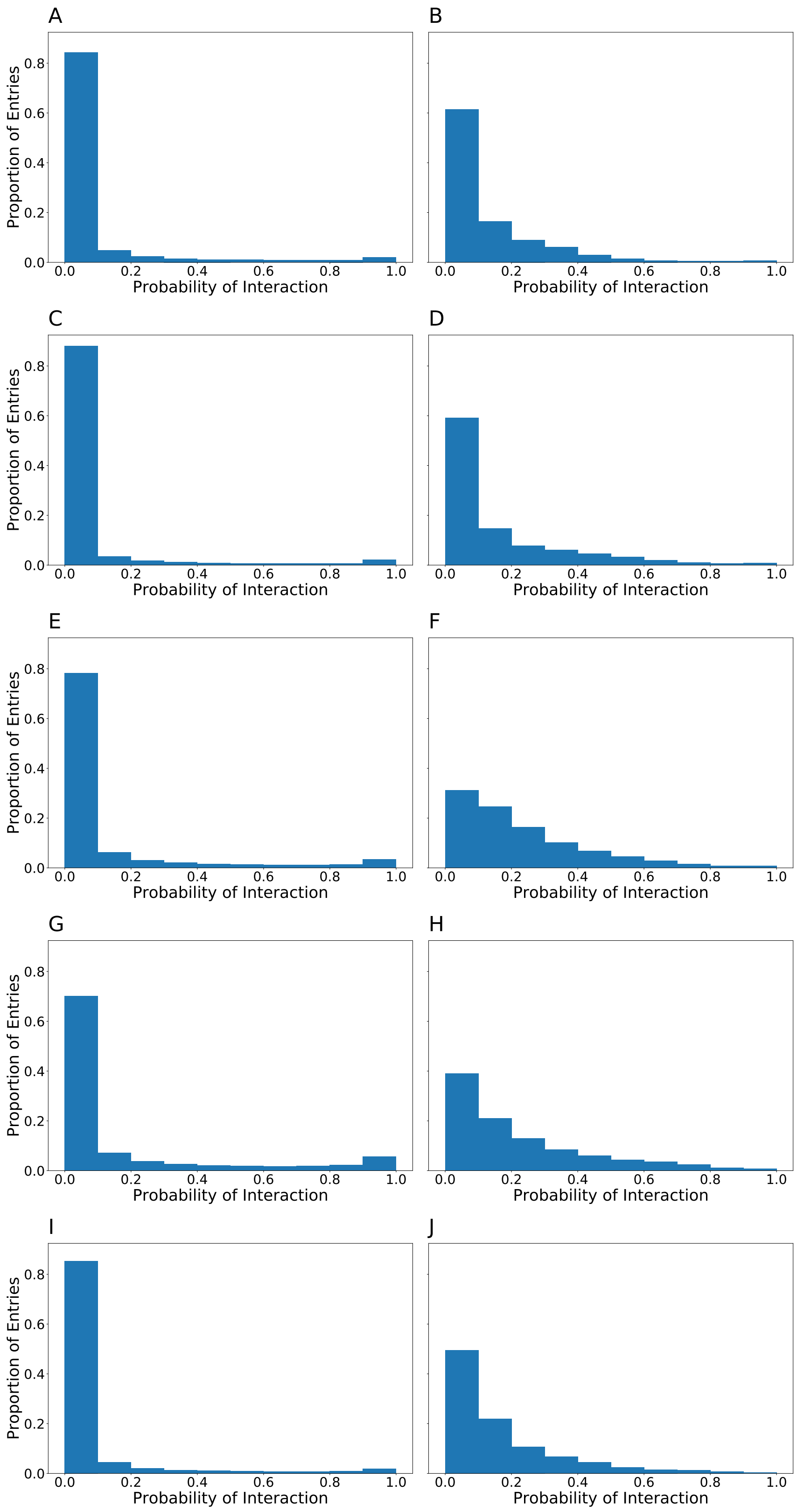


**Fig. S12.** Histogram of Added Information to Training Matrix for (a-b) Nuclear Receptor, (c-d) Ion Channel, (e-f) GPCR, (g-h) Kinase, and (i-j) Other Enzyme Datasets using (a, c, e, g, i) Logistic Regression and (b, d, f, h, j) Random Forest as Single Protein Models. This shows the predicted probabilities of interactions that are added to the training matrix by the appropriate single protein/ligand binding model. Most of the interactions are predicted to be inactive, but some are predicted to be active. We hypothesize that this additional data helps explain some of the improvement in performance.

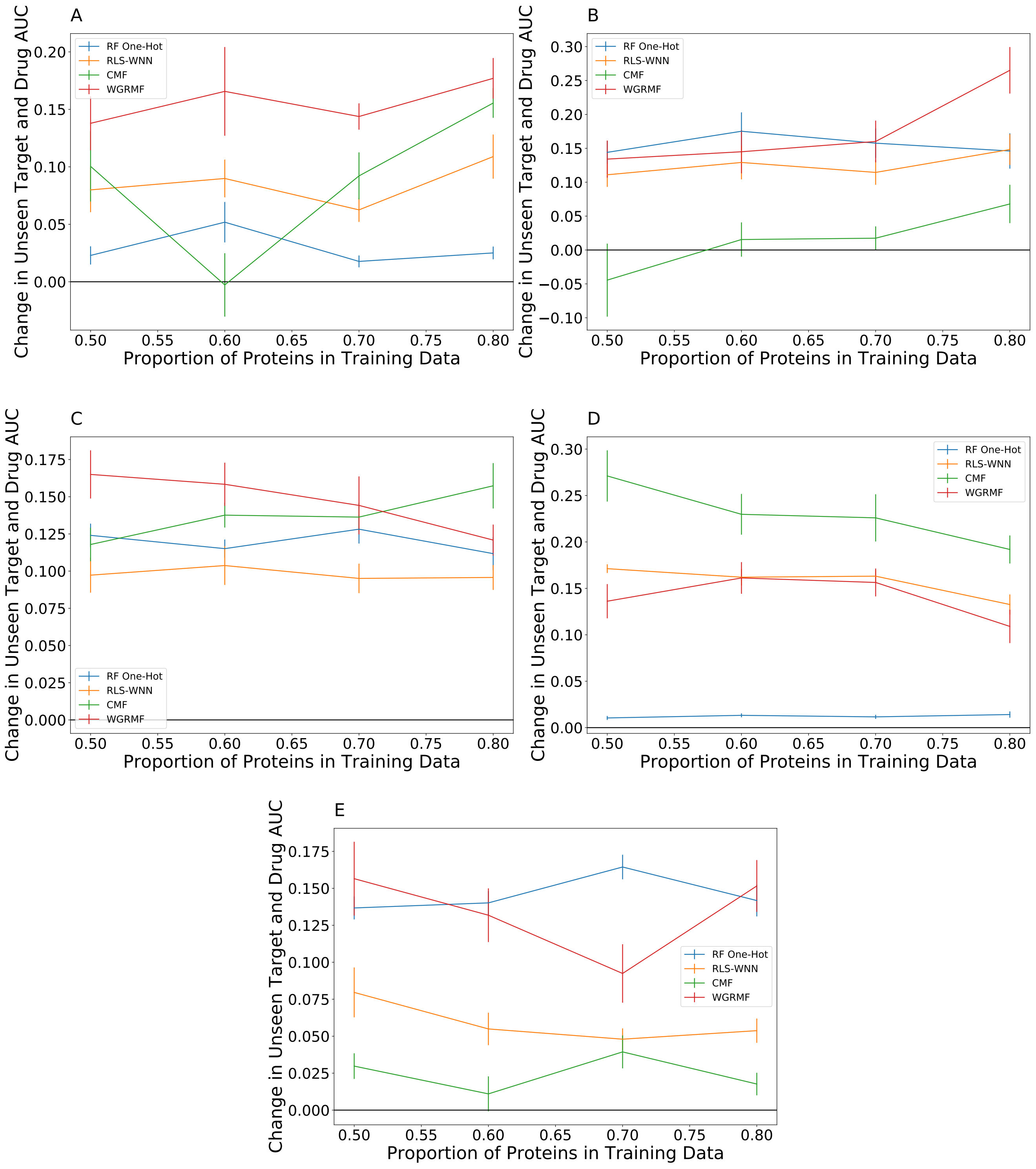


**Fig. S13.** Performance Dependence on Number of Proteins in Training Set for (a) Nuclear Receptor, (b) Ion Channel, (c) GPCR, (d) Kinase, and (e) Other Enzyme Datasets. Adding single protein/ligand binding models appears equally effective regardless of the number of proteins present in the training data.

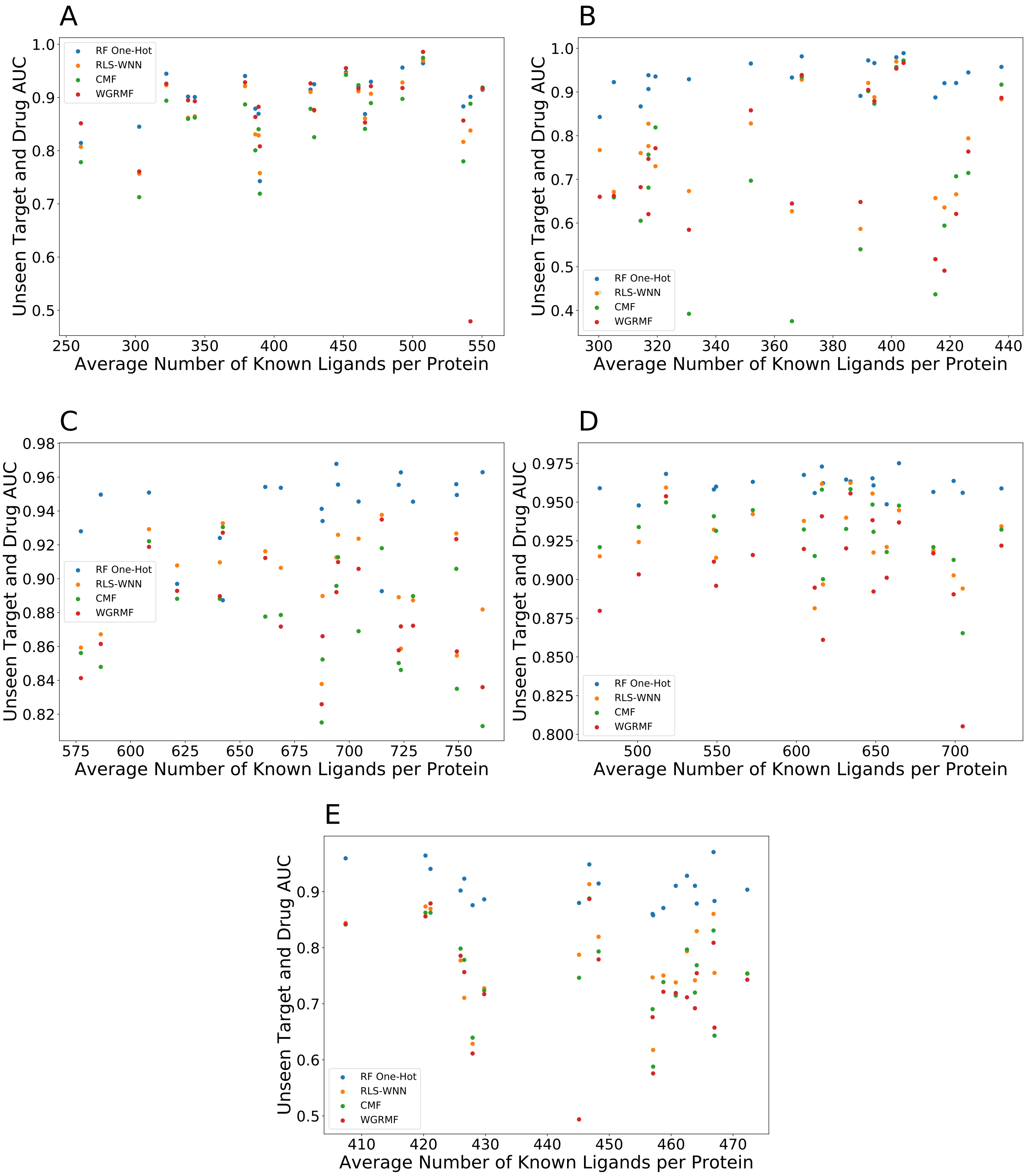


**Fig. S14.** Performance Dependence on Number of Known Active Ligands per Protein for (a) Nuclear Receptor, (b) Ion Channel, (c) GPCR, (d) Kinase, and (e) Other Enzyme Datasets. Our models' performance does not significantly depend on the average number of known ligands per protein.

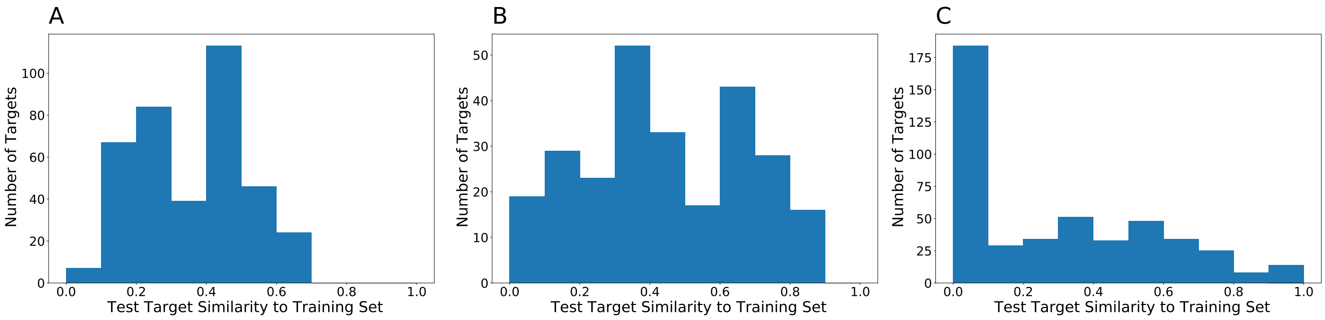


**Fig. S15.** Distribution of Target Similarity to Training Set for (A) the GPCR dataset, (B) the Kinase dataset, and (C) the Other Enzyme dataset. This histogram depicts the distribution of nearest-neighbor target similarity between all test targets and the training set across all $20$ trials for each dataset.

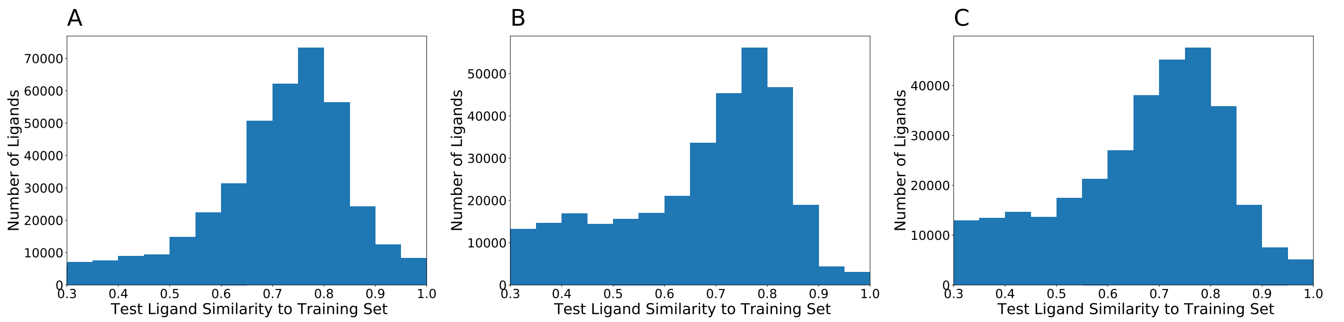


**Fig. S16.** Distribution of Ligand Similarity to Training Set for (A) the GPCR dataset, (B) the Kinase dataset, and (C) the Other Enzyme dataset. This histogram depicts the distribution of nearest-neighbor ligand similarity between all test ligands and the training set across all $20$ trials for each dataset.
