## Supplementary material for "Using Single Protein/Ligand Binding Models to Predict Active Ligands for Unseen Proteins": Code and dataset: AddSingleDTIClassifier.py

```
import numpy as np
from sklearn.linear_model import LogisticRegression
from sklearn.ensemble import RandomForestClassifier
import math

# This class adds a simple one protein/many ligand classifier to the one protein/many ligand problem. This returns probabilities, not classifications, so make sure the other classifier can appropriately accept probabilities in the input. If a proportion is given, it will only fill in the appropriate proportion of the training class using the model.

class AddSingleDTIClassifier:

    def __init__(self, otherclassifier, modelType="LR", proportion=1):
        self.modelType = modelType
        self.otherclassifier = otherclassifier
        self.proportion = proportion

    # submatrix method: takes in a training matrix, ligand list, list of training/test proteins and ligands, and outputs a completed test matrix
    def fit_predict_submatrix(self, trainmat, trainmatlog, trainmatrf, ligands, codes, trainprotvals, testprotvals, trainligvals, testligvals, X_trainonehotseq, X_testonehotseq, drugsim):
        if self.proportion == 1:
            if self.modelType == 'LR':
                trainmat = np.copy(trainmatlog)
            elif self.modelType == 'RF':
                trainmat = np.copy(trainmatrf)
            else:
                raise NotImplementedError
        else:
            numrows = math.floor(len(trainprotvals)*self.proportion)
            trainmat = np.copy(trainmat)
            if self.modelType == 'LR':
                trainmat[trainprotvals[:numrows], :] = trainmatlog[trainprotvals[:numrows], :]
            elif self.modelType == 'RF':
                trainmat[trainprotvals[:numrows], :] = trainmatrf[trainprotvals[:numrows], :]
            else:
                raise NotImplementedError

        # run the other classifier
        trainmat = self.otherclassifier.fit_predict_submatrix(trainmat, trainmatlog, trainmatrf, ligands, codes, trainprotvals, testprotvals, trainligvals, testligvals, X_trainonehotseq, X_testonehotseq, drugsim)
        return trainmat
```
