## Supplementary material for "Using Single Protein/Ligand Binding Models to Predict Active Ligands for Unseen Proteins": Code and dataset: AnalyzeDTIClassifier.py

```
import numpy as np
from rdkit import Chem
from rdkit.Chem import AllChem as Chem
import csv
from sklearn.model_selection import train_test_split
import matplotlib
matplotlib.use('agg')
import matplotlib.pyplot as plt
import math
import bisect
from itertools import product, chain
import sys
from scipy import sparse
from scipy.spatial.distance import pdist, cdist, squareform
from sklearn.metrics import roc_curve, roc_auc_score
from sklearn.linear_model import LogisticRegression
from sklearn.ensemble import RandomForestClassifier

# This script contains a number of helper methods used to compute AUCs for classifiers for the multiple protein/multiple ligand (i.e. drug/target interaction) problem. See the specific method for details.

percent_var = 0.2
num_decoys = 500

# helper method to do a submatrix split for a given DTI matrix
# splits the proteins and then does a split for both active and inactive ligands
def submatrix_split(testmat, codes, ligandsStr, decoysStr, activevals, inactivevals, percent_split, percent_split_ligand, trainprotvals=None, testprotvals=None, trainproteins=None, testproteins=None):
    trainmat = np.copy(testmat)
    if trainprotvals is None:
        trainprotvals, testprotvals, trainproteins, testproteins = train_test_split(range(len(codes)), codes, test_size = percent_split)

    # make sure we split out some active ligands and decoys for each protein
    trainactivevals = [[] for code in codes]
    testactivevals = [[] for code in codes]
    traininactivevals = [[] for code in codes]
    testinactivevals = [[] for code in codes]
    testactivevalsall = set()
    for i, code in enumerate(codes):
        trainactivevals[i], testactivevals[i] = train_test_split(activevals[i], test_size = percent_split_ligand)
        traininactivevals[i], testinactivevals[i] = train_test_split(inactivevals[i], test_size = percent_split_ligand)
        testactivevalsall |= set(testactivevals[i])
        testactivevalsall |= set(testinactivevals[i])
    trainactivevalsall = set(range(len(ligandsStr))) - testactivevalsall

    # do the overall decoy split
    traindecoyvals, testdecoyvals = train_test_split(range(len(ligandsStr), len(ligandsStr) + len(decoysStr)), test_size = percent_split_ligand)
    
    # combine everything together
    trainligvals = list(trainactivevalsall) + traindecoyvals
    testligvals = list(testactivevalsall) + testdecoyvals
    for prot, lig in product(testprotvals, range(len(ligandsStr) + len(decoysStr))):
        trainmat[prot][lig] = -1
    for prot, lig in product(range(len(codes)), testligvals):
        trainmat[prot][lig] = -1
   
    return trainmat, trainprotvals, testprotvals, trainproteins, testproteins, trainactivevalsall, testactivevalsall, traindecoyvals, testdecoyvals, trainligvals, testligvals

# helper method to generate train and test one-hot fingerprints
def generateOneHot(trainprotvals, testprotvals, trainactivevalsall, testactivevalsall, traindecoyvals, testdecoyvals, codes, ligandsStr, decoysStr, ligands):
    X_trainonehotseq = None
    X_testonehotseq = None
    for i in trainprotvals:
        # get all the fingerprints
        code = codes[i]
        X_traincurrentseq = np.loadtxt('/OneHot_Fingerprint_Sequence/CHEMBL' + code)

        # identify which ones need to be in test and train
        ligandcurrent = X_traincurrentseq[:, 20:-1]
        ligandcurrent = [''.join([str(val)[0] for val in ligand]) for ligand in ligandcurrent]
        todelete = []
        indices = []
        for k in testactivevalsall:
            curligand = ligandsStr[k]
            index = bisect.bisect(ligandcurrent, curligand)-1
            if ligandcurrent[index] == curligand:
                todelete += [index]
                indices += [k]
        tokeep = []
        for k in trainactivevalsall:
            curligand = ligandsStr[k]
            index = bisect.bisect(ligandcurrent, curligand)-1
            if ligandcurrent[index] == curligand:
                tokeep += [index]

        # split them into train and test
        X_testcurrentseq = np.hstack((np.tile([i], (len(indices),1)), np.array([indices]).T, X_traincurrentseq[todelete, :-1]))
        X_traincurrentseq = X_traincurrentseq[tokeep,:]

        # convert to sparse matrix
        seqfeatures = X_traincurrentseq[0, :20]
        X_traincurrentseq = sparse.lil_matrix(X_traincurrentseq)
        X_testcurrentseq = sparse.lil_matrix(X_testcurrentseq)

        # get the decoys and generate a matrix for them too
        X_traindecoyseq = np.hstack((np.tile(seqfeatures, (len(traindecoyvals),1)), ligands[traindecoyvals,:], np.tile([0], (len(traindecoyvals),1))))
        X_testdecoyseq = np.hstack((np.tile([i], (len(testdecoyvals),1)), np.array([testdecoyvals]).T, np.tile(seqfeatures, (len(testdecoyvals),1)), ligands[testdecoyvals,:]))

        # convert to sparse matrix
        X_traindecoyseq = sparse.lil_matrix(X_traindecoyseq)
        X_testdecoyseq = sparse.lil_matrix(X_testdecoyseq)

        # stack everything together
        X_traincurrentseq = sparse.vstack((X_traincurrentseq, X_traindecoyseq))
        X_testcurrentseq = sparse.vstack((X_testcurrentseq, X_testdecoyseq))

        # append to the overall answer
        if X_trainonehotseq is None:
            X_trainonehotseq = X_traincurrentseq
            X_testonehotseq = X_testcurrentseq
        else:
            X_trainonehotseq = sparse.vstack((X_trainonehotseq, X_traincurrentseq))
            X_testonehotseq = sparse.vstack((X_testonehotseq, X_testcurrentseq))

    # get the test proteins too
    print("Generating Test One-Hot Fingerprints")
    sys.stdout.flush()
    for i in testprotvals:
        # get all the fingerprints
        code = codes[i]
        X_testcurrentseq = np.loadtxt('/OneHot_Fingerprint_Sequence/CHEMBL' + code)[:, :-1]
        seqfeatures = X_testcurrentseq[0,:20]

        # identify indices and add them in
        ligandcurrent = X_testcurrentseq[:, 20:]
        ligandcurrent = [''.join([str(val)[0] for val in ligand]) for ligand in ligandcurrent]
        indices = []
        for ligand in ligandcurrent:
            index = bisect.bisect(ligandsStr, ligand)-1
            indices += [index]
        X_testcurrentseq = np.hstack((np.tile([i], (len(indices),1)), np.array([indices]).T, X_testcurrentseq))

        # convert to sparse matrix
        X_testcurrentseq = sparse.lil_matrix(X_testcurrentseq)

        # get decoys
        X_testdecoyseq = np.hstack((np.tile([i], (len(decoysStr),1)), np.array([range(len(ligandsStr),len(ligandsStr)+len(decoysStr))]).T, np.tile(seqfeatures, (len(decoysStr),1)), ligands[range(len(ligandsStr),len(ligandsStr)+len(decoysStr)),:]))
        X_testdecoyseq = sparse.lil_matrix(X_testdecoyseq)

        # stack everything together
        X_testcurrentseq = sparse.vstack((X_testcurrentseq, X_testdecoyseq))

        # add to the old matrix
        X_testonehotseq = sparse.vstack((X_testonehotseq, X_testcurrentseq))

    return X_trainonehotseq, X_testonehotseq

# helper method to fill in the training matrix with results from LR and RF
def fill_matrix(trainmat, ligands, codes, trainprotvals):
    trainmatlog = np.copy(trainmat)
    trainmatrf = np.copy(trainmat)
    # fill in the rows of the training matrix
    for i in trainprotvals:
        log = LogisticRegression(C=1, n_jobs=-1)
        rf = RandomForestClassifier(n_estimators=100, max_depth=25, n_jobs=-1)
        
        # set up a sub-training matrix and sub-testing matrix and train
        actives = ligands[trainmat[i, :]==1, :]
        inactives = ligands[trainmat[i, :]==0, :]
        X_train = np.vstack((actives, inactives))
        y_train = np.concatenate((np.repeat(1, np.shape(actives)[0]), np.repeat(0, np.shape(inactives)[0])))
        log.fit(X_train, y_train)
        rf.fit(X_train, y_train)

        # fill in the unknowns
        indices = np.arange(np.shape(trainmat)[1])[trainmat[i,:]==-1]
        X_test = ligands[trainmat[i,:]==-1,:]
        trainmatlog[i,indices] = log.predict_proba(X_test)[:,1]
        trainmatrf[i,indices] = rf.predict_proba(X_test)[:,1]

    return trainmatlog, trainmatrf

# quick helper method to compute AUC and ROC and store it
def aucCompute(trainmat, testmat, testmat_pred, true_posall, aucs, j, k, mean_fpr):
    indices = np.logical_and(trainmat==-1, testmat != -1)
    predvals = testmat_pred[indices]
    truevals = testmat[indices]
    fpr, tpr, thresholds = roc_curve(truevals, predvals)
    auc = roc_auc_score(truevals, predvals)
    aucs[k] += [auc]
    tpr_interp = np.interp(mean_fpr, fpr, tpr)
    tpr_interp[0] = 0
    tpr_interp[-1] = 1
    if j == 0:
        true_posall[k] = np.array([tpr_interp])
    else:
        true_posall[k] = np.vstack((true_posall[k], tpr_interp))
    print(auc)
    sys.stdout.flush()

# computes an ROC for the submatrix problem, i.e. a random set of proteins and ligands is omitted from the training matrix. Plots it as well if plot = True for the files in outputFiles. Else returns an AUC value.
# returns the false positive rate (+ error), false negative rate (+ error), and AUC (+ error), along with breakdowns for known drug/unknown target, unknown drug/known target, unknown drug/unknown target, and differences for each set of models run
# repeats computation repnum number of times to compute the error (repeats change the train/test set split)
# models should have a fit_predict_submatrix method for the submatrix case, i.e. taking just a training matrix and a list of ligands, test proteins, test ligands, and one-hot fingerprints, and filling in the training matrix
def SubmatrixROC(models, testmatnondecoy, codes, ligandsStr, drugsim, repnum=1, plot=False, plotinfo=[], hyperparams=False, percent_split_ligand=percent_var, targetsim=None, percent_split_prot=percent_var, splitFile=None, distanceFile=None, trainprotvals=None, testprotvals=None, trainproteins=None, testproteins=None):
    aucs = [[] for model in models]
    true_posall = [np.array([]) for model in models]
    aucuds = [[] for model in models]
    true_posudall = [np.array([]) for model in models]
    aucuts = [[] for model in models]
    true_posutall = [np.array([]) for model in models]
    aucuduts = [[] for model in models]
    true_posudutall = [np.array([]) for model in models]
    aucpertargets = [[[] for model in models] for i in range(math.ceil(percent_split_prot*np.shape(testmatnondecoy)[0]))]
    true_pospertargetsall =  [[np.array([]) for model in models] for i in range(math.ceil(percent_split_prot*np.shape(testmatnondecoy)[0]))]
    aucdiffs = [[[] for model in models] for model in models]
    aucuddiffs = [[[] for model in models] for model in models]
    aucutdiffs = [[[] for model in models] for model in models]
    aucudutdiffs = [[[] for model in models] for model in models]
    # test matrix of drug/target interactions - will have 1 for known active, 0 for known inactive, -1 for unknown
    activevals = [np.nonzero(testmatnondecoy[i,:] == 1)[0] for i in range(np.shape(testmatnondecoy)[0])]
    inactivevals = [np.nonzero(testmatnondecoy[i,:] == 0)[0] for i in range(np.shape(testmatnondecoy)[0])]
    # list of ligands by Morgan3 fingerprint
    ligandarr = np.array([list(map(int, ligand)) for ligand in ligandsStr])

    # repetition to get error bars
    for j in range(repnum):
        print("Beginning Repetition Number " + str(j))
        sys.stdout.flush()
        
        # generate decoys randomly
        print("Generating Decoys")
        sys.stdout.flush()
        ligandset = set(ligandsStr)
        decoyset = set()
        # read in the ChEMBL database
        with open('chembl_24_1_chemreps.txt') as f:
            reader = csv.reader(f.readlines(), delimiter='\t')
            d = list(reader)
        # add additional decoys
        notdone = True
        while notdone:
            smi = [item[1] for item in d[1:]]
            index = np.random.permutation(len(smi))
            ms = []
            for x in np.arange(num_decoys):
                if Chem.MolFromSmiles(smi[index[x]]) is not None:
                    ms.append(Chem.MolFromSmiles(smi[index[x]]))
                else:
                    ms.append(Chem.MolFromSmiles(smi[index[x]], sanitize=False))
                    print(x)
            try:
                decoyset = set([Chem.GetMorganFingerprintAsBitVect(y, 3).ToBitString() for y in ms])
                notdone = False
            except RuntimeError:
                continue
        decoyset -= set(ligandset)
        decoysStr = list(decoyset)
        decoysStr.sort()
        
        testmat = np.hstack((testmatnondecoy, np.zeros((len(codes), len(decoysStr)))))
        decoys = np.array([list(map(int, decoy)) for decoy in decoysStr])
        ligands = np.vstack((ligandarr, decoys))
       
        # Update the distance matrix
        print("Updating Distance Matrix")
        sys.stdout.flush()
        decoydecoysim = 1 - squareform(pdist(decoys, metric='jaccard'))
        decoytargetsim = 1 - cdist(ligandarr, decoys, metric='jaccard')
        drugsimfinal = np.vstack((np.hstack((drugsim, decoytargetsim)), np.hstack((decoytargetsim.T, decoydecoysim))))

        # now split the train/test matrices
        print("Splitting into Train/Test")
        sys.stdout.flush()
        trainmat, trainprotvals, testprotvals, trainproteins, testproteins, trainactivevalsall, testactivevalsall, traindecoyvals, testdecoyvals, trainligvals, testligvals = submatrix_split(testmat, codes, ligandsStr, decoysStr, activevals, inactivevals, percent_split_prot, percent_split_ligand, trainprotvals=trainprotvals, testprotvals=testprotvals, trainproteins=trainproteins, testproteins=testproteins)
        if distanceFile is not None:
            np.save(distanceFile, targetsim[np.ix_(trainprotvals, testprotvals)])
        if splitFile is not None:
            numligs = []
            for prot in trainprotvals:
                numligs += [np.shape(trainmat[prot, trainmat[prot, :] == 1])]
            np.save(splitFile, numligs)
        
        # prepare a training and test one-hot fingerprint matrix - the test matrix also begins with the protein/ligand index to speed up preparation of the answer for the models
        print("Generating One-Hot Fingerprints")
        sys.stdout.flush()
        X_trainonehotseq, X_testonehotseq = generateOneHot(trainprotvals, testprotvals, trainactivevalsall, testactivevalsall, traindecoyvals, testdecoyvals, codes, ligandsStr, decoysStr, ligands)  
        
        # tune hyperparameters as necessary
        print("Tuning Hyperparameters")
        sys.stdout.flush()
        bestmodels = models
        if hyperparams:
            # pull out the subset we care about
            subtestmat = trainmat[np.ix_(trainprotvals, trainligvals)]
            subcodes = [codes[protval] for protval in trainprotvals]
            subligandsStr = [ligandsStr[activeval] for activeval in trainactivevalsall]
            subdecoysStr = [decoysStr[decoyval-len(ligandsStr)] for decoyval in traindecoyvals]
            subligands = ligands[trainligvals, :]
            subactivevals = [np.nonzero(subtestmat[i,:len(subligandsStr)] == 1)[0] for i in range(np.shape(subtestmat)[0])]
            subinactivevals = [np.nonzero(subtestmat[i,:len(subligandsStr)] == 0)[0] for i in range(np.shape(subtestmat)[0])]
            
            # split into train/validation
            print("Splitting into Train/Validation")
            sys.stdout.flush()
            htrainmat, htrainprotvals, hvalprotvals, htrainproteins, hvalproteins, htrainactivevalsall, hvalactivevalsall, htraindecoyvals, hvaldecoyvals, htrainligvals, hvalligvals = submatrix_split(subtestmat, subcodes, subligandsStr, subdecoysStr, subactivevals, subinactivevals, percent_var/(1 - percent_var), percent_var/(1 - percent_var))

            # generate one-hot fingerprints
            print("Generating One-Hot Fingerprints")
            sys.stdout.flush()
            X_htrainonehotseq, X_htestonehotseq = generateOneHot(htrainprotvals, hvalprotvals, htrainactivevalsall, hvalactivevalsall, htraindecoyvals, hvaldecoyvals, subcodes, subligandsStr, subdecoysStr, subligands)

            # fill in the matrix with results from LR/RF
            print("Filling In Training Matrix")
            sys.stdout.flush()
            htrainmatlog, htrainmatrf = fill_matrix(htrainmat, subligands, subcodes, htrainprotvals)

            # run through all the models
            for k, modeloptions in enumerate(models):
                if len(modeloptions) == 1:
                    bestmodels[k] = modeloptions[0]
                else:  
                    print("Tuning Hyperparameters for Model " + str(k))
                    sys.stdout.flush()

                    mean_fpr = np.linspace(0, 1, 1000)
                    model_aucs = [[] for l in range(len(modeloptions))]
                    model_true_posall = [[] for l in range(len(modeloptions))]
                    for l in range(len(modeloptions)):
                        # fit the model
                        print("Evaluating Option Number " + str(l))
                        sys.stdout.flush()
                        testmat_pred = modeloptions[l].fit_predict_submatrix(htrainmat, htrainmatlog, htrainmatrf, subligands, subcodes, htrainprotvals, hvalprotvals, htrainligvals, hvalligvals, X_htrainonehotseq, X_htestonehotseq, drugsimfinal[np.ix_(trainligvals, trainligvals)])
                        
                        # compute the AUC and store it
                        aucCompute(htrainmat, subtestmat, testmat_pred, model_true_posall, model_aucs, 0, l, mean_fpr)
                    bestmodels[k] = modeloptions[np.argmax(model_aucs)]

        # fill in the matrix with results from LR/RF
        print("Filling In Training Matrix")
        sys.stdout.flush()
        trainmatlog, trainmatrf = fill_matrix(trainmat, ligands, codes, trainprotvals)

        mean_fpr = np.linspace(0, 1, 1000)
        testmatud = testmat[np.ix_(trainprotvals, testligvals)]
        trainmatud = trainmat[np.ix_(trainprotvals, testligvals)]
        testmatut = testmat[np.ix_(testprotvals, trainligvals)]
        trainmatut = trainmat[np.ix_(testprotvals, trainligvals)]
        testmatudut = testmat[np.ix_(testprotvals, testligvals)]
        trainmatudut = trainmat[np.ix_(testprotvals, testligvals)]
        # evaluate the models
        for k, model in enumerate(models):
            print("Evaluating Model Number " + str(k))
            sys.stdout.flush()

            # fit the model and get predictions
            testmat_pred = model.fit_predict_submatrix(trainmat, trainmatlog, trainmatrf, ligands, codes, trainprotvals, testprotvals, trainligvals, testligvals, X_trainonehotseq, X_testonehotseq, drugsimfinal)

            # compute the AUC and store it
            aucCompute(trainmat, testmat, testmat_pred, true_posall, aucs, j, k, mean_fpr)

            # do the same for unknown drug/known target
            testmatud_pred = testmat_pred[np.ix_(trainprotvals, testligvals)]
            aucCompute(trainmatud, testmatud, testmatud_pred, true_posudall, aucuds, j, k, mean_fpr)

            # do the same for known drug/unknown target
            testmatut_pred = testmat_pred[np.ix_(testprotvals, trainligvals)]
            aucCompute(trainmatut, testmatut, testmatut_pred, true_posutall, aucuts, j, k, mean_fpr)
            
            # do the same for unknown drug/unknown target
            testmatudut_pred = testmat_pred[np.ix_(testprotvals, testligvals)]
            aucCompute(trainmatudut, testmatudut, testmatudut_pred, true_posudutall, aucuduts, j, k, mean_fpr)

            # do the same for each unknown target
            for l, prot in enumerate(testprotvals):
                testmatprot_pred = testmat_pred[np.ix_([prot], testligvals)]
                testmatprot = testmat[np.ix_([prot], testligvals)]
                trainmatprot = trainmat[np.ix_([prot], testligvals)]
                aucCompute(trainmatprot, testmatprot, testmatprot_pred, true_pospertargetsall[l], aucpertargets[l], j, k, mean_fpr)

        for k in range(len(models)):
            for l in range(len(models)):
                aucdiffs[k][l] += [aucs[k][-1] - aucs[l][-1]]
                aucutdiffs[k][l] += [aucuts[k][-1] - aucuts[l][-1]]
                aucuddiffs[k][l] += [aucuds[k][-1] - aucuds[l][-1]]
                aucudutdiffs[k][l] += [aucuduts[k][-1] - aucuduts[l][-1]]
    
    aucs = np.array(aucs)
    true_posall = np.array(true_posall)
    aucuts = np.array(aucuts)
    true_posutall = np.array(true_posutall)
    aucuds = np.array(aucuds)
    true_posudall = np.array(true_posudall)
    aucuduts = np.array(aucuduts)
    true_posudutall = np.array(true_posudutall)
    aucpertargets = np.array(aucpertargets)
    true_pospertargetsall = np.array(true_pospertargetsall)
    aucdiffs = np.array(aucdiffs)
    aucutdiffs = np.array(aucutdiffs)
    aucuddiffs = np.array(aucuddiffs)
    aucudutdiffs = np.array(aucudutdiffs)

    # compute means and standard deviations as necessary
    aucmean = np.mean(aucs, axis=1)
    aucstd = np.std(aucs, axis=1)/math.sqrt(repnum)
    true_posmean = np.mean(true_posall, axis=1)
    true_posstd = np.std(true_posall, axis=1)/math.sqrt(repnum)
    aucudmean = np.mean(aucuds, axis=1)
    aucudstd = np.std(aucuds, axis=1)/math.sqrt(repnum)
    true_posudmean = np.mean(true_posudall, axis=1)
    true_posudstd = np.std(true_posudall, axis=1)/math.sqrt(repnum)
    aucutmean = np.mean(aucuts, axis=1)
    aucutstd = np.std(aucuts, axis=1)/math.sqrt(repnum)
    true_posutmean = np.mean(true_posutall, axis=1)
    true_posutstd = np.std(true_posutall, axis=1)/math.sqrt(repnum)
    aucudutmean = np.mean(aucuduts, axis=1)
    aucudutstd = np.std(aucuduts, axis=1)/math.sqrt(repnum)
    true_posudutmean = np.mean(true_posudutall, axis=1)
    true_posudutstd = np.std(true_posudutall, axis=1)/math.sqrt(repnum)
    aucdiffmean = np.mean(aucdiffs, axis=2)
    aucdiffstd = np.std(aucdiffs, axis=2)/math.sqrt(repnum)
    aucutdiffmean = np.mean(aucutdiffs, axis=2)
    aucutdiffstd = np.std(aucutdiffs, axis=2)/math.sqrt(repnum)
    aucuddiffmean = np.mean(aucuddiffs, axis=2)
    aucuddiffstd = np.std(aucuddiffs, axis=2)/math.sqrt(repnum)
    aucudutdiffmean = np.mean(aucudutdiffs, axis=2)
    aucudutdiffstd = np.std(aucudutdiffs, axis=2)/math.sqrt(repnum)
        
    # plot the ROC curve as necessary
    if plot:
        for plotmodels, plotlabels, plotfiles in plotinfo:
            fig, ax = plt.subplots()
            for i, k in enumerate(plotmodels):
                ax.errorbar(mean_fpr, true_posmean[k], yerr=true_posstd[k], label=plotlabels[i])
            ax.set_xlim([0, 1])
            ax.set_ylim([0, 1])
            ax.set_xlabel("False positive")
            ax.set_ylabel("True positive")
            plt.legend()
            plt.savefig(plotfiles + "AllROC.jpg")

            fig, ax = plt.subplots()
            for i, k in enumerate(plotmodels):
                ax.errorbar(mean_fpr, true_posutmean[k], yerr=true_posutstd[k], label=plotlabels[i])
            ax.set_xlim([0, 1])
            ax.set_ylim([0, 1])
            ax.set_xlabel("False positive")
            ax.set_ylabel("True positive")
            plt.legend()
            plt.savefig(plotfiles + "UnknownTargetROC.jpg")

            fig, ax = plt.subplots()
            for i, k in enumerate(plotmodels):
                ax.errorbar(mean_fpr, true_posudmean[k], yerr=true_posudstd[k], label=plotlabels[i])
            ax.set_xlim([0, 1])
            ax.set_ylim([0, 1])
            ax.set_xlabel("False positive")
            ax.set_ylabel("True positive")
            plt.legend()
            plt.savefig(plotfiles + "UnknownDrugROC.jpg")
            
            fig, ax = plt.subplots()
            for i, k in enumerate(plotmodels):
                ax.errorbar(mean_fpr, true_posudutmean[k],yerr=true_posudutstd[k], label=plotlabels[i])
            ax.set_xlim([0, 1])
            ax.set_ylim([0, 1])
            ax.set_xlabel("False positive")
            ax.set_ylabel("True positive")
            plt.legend()
            plt.savefig(plotfiles + "BothUnknownROC.jpg")
    
    return [true_posmean, true_posstd, aucmean, aucstd, true_posutmean, true_posutstd, aucutmean, aucutstd, true_posudmean, true_posudstd, aucudmean, aucudstd, true_posudutmean, true_posudutstd, aucudutmean, aucudutstd, aucdiffmean, aucdiffstd, aucutdiffmean, aucutdiffstd, aucuddiffmean, aucuddiffstd, aucudutdiffmean, aucudutdiffstd, aucpertargets, true_pospertargetsall]
```
