## Supplementary material for "Using Single Protein/Ligand Binding Models to Predict Active Ligands for Unseen Proteins": Code and dataset: CMFDTIClassifier.py

```
import numpy as np
from numpy import linalg
from scipy.sparse.linalg import svds
from scipy import sparse
import pickle

# This class implements the CMF classifier as described in Zheng 2013. It attempts to factor the drug/target interaction matrix trainmat as AB^T with a bunch of regularization penalties characterized by lambda_l (overall matrix norm), lambda_d (deviation from the drug similarity), and lambda_t (deviation from target similarity). k is the number of latent features (must be < than the number of targets).

class CMFDTIClassifier:

    def __init__(self, similarity, similarityCodes, k, lambda_l, lambda_d, lambda_t):
        self.similarity = similarity
        self.similarityCodes = similarityCodes
        self.k = k
        self.lambda_l = lambda_l
        self.lambda_d = lambda_d
        self.lambda_t = lambda_t

    # submatrix method: takes in a training matrix, ligand list, list of training/test proteins and ligands, and outputs a completed test matrix
    def fit_predict_submatrix(self, trainmat, trainmatlog, trainmatrf, ligands, codes, trainprotvals, testprotvals, trainligvals, testligvals, X_trainonehotseq, X_testonehotseq, drugsim):
        self.k = min(self.k, np.shape(trainmat)[0]-1)
        trainmat = np.copy(trainmat)
        W = (trainmat != -1)
        trainmat[trainmat == -1] = 0
        targetsim = np.zeros((len(codes), len(codes)))
        for i, code in enumerate(codes):
            for j, othercode in enumerate(codes):
                targetsim[i][j] = self.similarity[self.similarityCodes.index(code)][self.similarityCodes.index(othercode)]
        losses = []

        # initialize latent feature matrices and weight matrix
        trainmatsparse = sparse.lil_matrix(trainmat)
        u, s, vh = svds(trainmatsparse, k=self.k)
        v = vh.T
        A = u*np.sqrt(s)
        B = v*np.sqrt(s)
        loss = np.linalg.norm(np.multiply(W, trainmat - np.dot(A, B.T))) + self.lambda_l*(np.linalg.norm(A) + np.linalg.norm(B)) + self.lambda_d*np.linalg.norm(drugsim - np.dot(B, B.T)) + self.lambda_t*np.linalg.norm(targetsim - np.dot(A, A.T))
        losses += [loss]

        # run the iterations
        lambda_d_Sd = self.lambda_d*drugsim
        lambda_t_St = self.lambda_t*targetsim
        lambda_l_ik = self.lambda_l*np.identity(self.k)
        H = np.multiply(W, trainmat)

        # see the Ezzat paper's Github for the code I reimplemented here
        notdone = True
        count = 0
        while notdone and count < 20:
            count += 1
            A_old = A
            B_old = B
            numA = np.dot(H, B) + np.dot(lambda_t_St,A_old)
            ltAA = lambda_l_ik + self.lambda_t*np.dot(A_old.T, A_old)
            for a in range(np.shape(A)[0]):
                A[a,:] = linalg.solve((np.dot(B.T*W[a,:], B) + ltAA).T, (numA[a,:]).T).T
            numB = np.dot(H.T, A) + np.dot(lambda_d_Sd,B_old)
            ldBB = lambda_l_ik + self.lambda_d*np.dot(B_old.T, B_old)
            for b in range(np.shape(B)[0]):
                B[b,:] = linalg.solve((np.dot(A.T*W[:,b], A) + ldBB).T, (numB[b,:]).T).T
            loss = np.linalg.norm(np.multiply(W, trainmat - np.dot(A, B.T))) + self.lambda_l*(np.linalg.norm(A) + np.linalg.norm(B)) + self.lambda_d*np.linalg.norm(drugsim - np.dot(B, B.T)) + self.lambda_t*np.linalg.norm(targetsim - np.dot(A, A.T))
            losses += [loss]
            if count%2 == 0 and losses[-1] > losses[-3] - 10:
                notdone = False

        # return the answer
        return np.dot(A, B.T)
```
