## Supplementary material for "Using Single Protein/Ligand Binding Models to Predict Active Ligands for Unseen Proteins": Code and dataset: CompileLigandList.py

```
import numpy as np
from rdkit import Chem
from rdkit.Chem import AllChem as Chem
import csv
import bisect
from collections import Counter
from scipy.spatial.distance import pdist, squareform
import sys

# This script takes a list of CHEMBL codes and compiles a sorted list (with no repeats) of all the active/inactive ligands and their fingerprints. It also computes the one-hot fingerprints for a given dataset and the Tanimoto fingerprint similarity matrix. Due to the extensive computation, this code can take a while to run.
# This code requires the Active_Fingerprint_Data/ and Inactive_Fingerprint_Data/ to be already generated by ParseCHEMBLAll.py and PubChemDecoys.py. It also requires the DUD-E decoy set to be already manually downloaded. 
# It stores the resulting feature sets in OneHot_Fingerprint_Sequence/. It also stores similarity matrices in DrugSim_*.npy, DTI matrices in DTI_matrix_*.in, and the ligand lists in LigandList_*.txt. Note that the drug similarity matrices can be very large (30+ GB).

dataset = sys.argv[1]

codes = []
with open('codes_' + dataset + '.txt') as f:
    for line in f:
        codes += [line[:-1]]

# get all the actives and inactives and DUDE decoys
ligandset = set()
numactives = 0
numinactives = 0
todelete = []
for code in codes:
    print(len(ligandset))
    sys.stdout.flush()
    worked = True
    try:
        actives = np.loadtxt("Active_Fingerprint_Data/CHEMBL" + code)
        if len(np.shape(actives)) == 1 or len(actives) < 20:
            print("CODE " + code)
            worked = False
            todelete += [code]
            sys.stdout.flush()
        else:
            actives = actives[:,:-1]
            numactives += np.shape(actives)[0]
            for active in actives:
                active = ''.join(str(int(e))[0] for e in active.tolist())
                if active not in ligandset:
                    ligandset.add(active)
    except IOError:
        print("CODE " + code)
        worked = False
        todelete += [code]
        sys.stdout.flush()
    if worked:
        inactives = []
        try:
            inactives = np.loadtxt("Inactive_Fingerprint_Data/CHEMBL" + code)
            if len(inactives) > 2000:
                inactives = inactives[:2000]
        except IOError:
            continue
        if len(np.shape(inactives)) > 1:
            numinactives += np.shape(inactives)[0]
            for inactive in inactives:
                inactive = ''.join(str(int(e))[0] for e in inactive.tolist())
                if inactive not in ligandset:
                    ligandset.add(inactive)
        decoyssmiles = []
        try:
            with open('DUDE_Decoys/CHEMBL' + code) as f:
                for line in f:
                    lineSplit = line.split(" ")
                    decoyssmiles += [lineSplit[0]]
            numinactives += min(len(decoyssmiles), np.shape(actives)[0])
            for decoysmi in decoyssmiles[:np.shape(actives)[0]]:
                if Chem.MolFromSmiles(decoysmi) is not None:
                    mol = Chem.MolFromSmiles(decoysmi)
                    try:
                        decoy = Chem.GetMorganFingerprintAsBitVect(mol,3).ToBitString()
                        if decoy not in ligandset:
                            ligandset.add(decoy)
                    except RuntimeError:
                        continue
        except IOError:
            continue

codes = [code for code in codes if code not in todelete]
ligandset = list(ligandset)
ligandset.sort()
ligands = np.array([list(map(int, ligand)) for ligand in ligandset])
DTImat = -np.ones((len(codes), len(ligandset)))

# label columns as active and inactive and prepare the DTI matrix
for i, code in enumerate(codes):
    print(i)
    sys.stdout.flush()
    actives = np.loadtxt("Active_Fingerprint_Data/CHEMBL" + str(code))
    actives = actives[:,:-1]
    for active in actives:
        activeStr = ''.join(str(int(e))[0] for e in active.tolist())
        index = bisect.bisect(ligandset, activeStr)-1
        DTImat[i][index] = 1
    try:
        inactives = np.loadtxt("Inactive_Fingerprint_Data/CHEMBL" + str(code))
        if len(inactives) > 2000:
            inactives = inactives[:2000]
    except IOError:
        inactives = np.array([])
    if len(np.shape(inactives)) > 1:
        for inactive in inactives:
            inactiveStr = ''.join(str(int(e))[0] for e in inactive.tolist())
            index = bisect.bisect(ligandset, inactiveStr)-1
            DTImat[i][index] = 0
    try:
        decoyssmiles = []
        with open('DUDE_Decoys/CHEMBL' + code) as f:
            for line in f:
                lineSplit = line.split(" ")
                decoyssmiles += [lineSplit[0]]
        for decoysmi in decoyssmiles[:np.shape(actives)[0]]:
            if Chem.MolFromSmiles(decoysmi) is not None:
                mol = Chem.MolFromSmiles(decoysmi)
                try:
                    decoy = Chem.GetMorganFingerprintAsBitVect(mol,3).ToBitString()
                    index = bisect.bisect(ligandset, decoy)-1
                    DTImat[i][index] = 0
                except RuntimeError:
                    continue
    except IOError:
        continue

np.savetxt('DTI_matrix_' + dataset + '.in', DTImat)

print(numactives, numinactives)
sys.stdout.flush()

# prepare one-hot fingerprints
for i, code in enumerate(codes):
    ligandvals = np.nonzero(DTImat[i, :]!=-1)[0]
    aa_list = ['A', 'C', 'D', 'E', 'F', 'G', 'H', 'I', 'K', 'L', 'M', 'N', 'P', 'Q', 'R', 'S', 'T', 'V', 'W', 'Y']
    with open("Real_Sequences/CHEMBL" + code) as f:
        line = f.readline()
        realseq = f.readline()
    countreal = Counter(realseq)
    realfeatures = np.array([countreal[char] for char in aa_list])
    realfeaturesall = np.array([])
    for val in ligandvals:
        realligfeatures = np.concatenate((realfeatures, ligands[val, :], np.array([DTImat[i, val]])))
        if len(realfeaturesall) == 0:
            realfeaturesall = np.array([realligfeatures])
        else:
            realfeaturesall = np.vstack((realfeaturesall, realligfeatures))
    np.savetxt('OneHot_Fingerprint_Sequence/CHEMBL' + code, realfeaturesall)

# get similarities and store them
drugsim = 1 - squareform(pdist(ligands, metric='jaccard'))
np.save('DrugSim_' + dataset + '.npy', drugsim)

# write out
with open('LigandList_' + dataset + '.txt', 'w') as f:
    for ligand in ligandset:
        f.write(ligand + '\n')
```
