## Supplementary material for "Using Single Protein/Ligand Binding Models to Predict Active Ligands for Unseen Proteins": Code and dataset: DTIModelsCompare.py

```
import numpy as np
import AnalyzeDTIClassifier
from WeightedNNDTIClassifier import WeightedNNDTIClassifier
from OneHotDTIClassifier import OneHotDTIClassifier
from WeightedProfileDTIClassifier import WeightedProfileDTIClassifier
from RLSWNNDTIClassifier import RLSWNNDTIClassifier
from CMFDTIClassifier import CMFDTIClassifier
from WGRMFDTIClassifier import WGRMFDTIClassifier
from AddSingleDTIClassifier import AddSingleDTIClassifier
from RLSWNNSingleDTIClassifier import RLSWNNSingleDTIClassifier
from OneHotSingleDTIClassifier import OneHotSingleDTIClassifier
import sys
import os
import subprocess

# This script runs a bunch of DTI models and produces the results. You can specify which models to run and what run number is used for storage. We assume the similarity matrices and the DTI matrix have already been computed prior to running this script.
# All results are stored in Analysis/DTI/<dataset>run<runno>, which is overwritten if necessary.

dataset = sys.argv[1]
runno = int(sys.argv[2])
os.environ['OMP_NUM_THREADS'] = '32'

# create and clean out output directory
subprocess.call("mkdir Analysis/DTI/" + dataset + "run" + str(runno) + "/", shell=True)
subprocess.call("rm Analysis/DTI/" + dataset + "run" + str(runno) + "/*", shell=True)
outfolder = "Analysis/DTI/" + dataset + "run" + str(runno) + "/"

# read in similarity matrices and the list of codes
sim = np.loadtxt('Similarity/Similarity_Sequence_' + dataset + '.out')
drugsim = np.load('DrugSim_' + dataset + '.npy')
DTImatrix = np.loadtxt('DTI_matrix_' + dataset + '.in')
codes = []
with open('codes_' + dataset + '.txt') as f:
    for line in f:
        codes += [line[:-1]]
ligands = []
with open('LigandList_' + dataset + '.txt') as f:
    for line in f:
        ligands += [line[:-1]]

# models
WeightedProfile = [WeightedProfileDTIClassifier(similarity=sim, similarityCodes=codes)]
WeightedNNLR = [WeightedNNDTIClassifier(similarity=sim, similarityCodes=codes, modelType="LR")]
WeightedNNRF = [WeightedNNDTIClassifier(similarity=sim, similarityCodes=codes, modelType="RF")]

RFOne = []
RFOneAddLog = []
RFOneAddRF = []
for num_trees in np.arange(50, 200, step=50):
    for max_depth in np.arange(15, 35, step=5):
        RFOne += [OneHotDTIClassifier(sequence='Seq', model='RF', hyperparams=[num_trees,max_depth])]
        RFOneAddLog += [OneHotSingleDTIClassifier(sequence='Seq', model='RF', hyperparams=[num_trees,max_depth], singlemodel='LR')]
        RFOneAddRF += [OneHotSingleDTIClassifier(sequence='Seq', model='RF', hyperparams=[num_trees,max_depth], singlemodel='RF')]

RLSWNN = []
RLSWNNSingleLog = []
RLSWNNSingleRF = []
hyperparams = [[0.25, 0.1, 0.4], [0.5, 0.9, 0.4], [2, 0.6, 0.1], [0.5, 1, 0.6]]
for val in hyperparams:
    RLSWNN += [RLSWNNDTIClassifier(similarity=sim, similarityCodes=codes, eta=val[2], alpha=val[1], sigma=val[0], k=100)]
hyperparams = [[0.5, 0.5, 0.6], [0.5, 1, 0.7], [0.25, 1, 0.7]]
for val in hyperparams:
    RLSWNNSingleLog += [RLSWNNSingleDTIClassifier(similarity=sim, similarityCodes=codes, eta=val[2], alpha=val[1], sigma=val[0], modelType="LR")]
    RLSWNNSingleRF += [RLSWNNSingleDTIClassifier(similarity=sim, similarityCodes=codes, eta=val[2], alpha=val[1], sigma=val[0], modelType="RF")]

CMF = []
CMFAddLog = []
CMFAddRF = []
hyperparams = [[1, 4, 32], [2, 8, 0.125], [4, 32, 0.25], [2, 64, 0.125], [0.25, 0.25, 32], [1, 0.0625, 2]]
for val in hyperparams:
    CMF += [CMFDTIClassifier(similarity=sim, similarityCodes=codes, k=30, lambda_l=val[0], lambda_d=val[1], lambda_t=val[2])]
hyperparams = [[1, 4, 32], [0.25, 0.25, 32], [0.5, 0.0625, 64], [0.5, 0.0625, 32]]
for val in hyperparams:
    CMFAddLog += [AddSingleDTIClassifier(modelType="LR", otherclassifier=CMFDTIClassifier(similarity=sim, similarityCodes=codes, k=30, lambda_l=val[0], lambda_d=0, lambda_t=val[2]))]
    CMFAddRF += [AddSingleDTIClassifier(modelType="RF", otherclassifier=CMFDTIClassifier(similarity=sim, similarityCodes=codes, k=30, lambda_l=val[0], lambda_d=0, lambda_t=val[2]))]

WGRMF = []
WGRMFAddLog = []
WGRMFAddRF = []
hyperparams = [[0.0625, 0.05, 0.1, 2], [0.25, 0.2, 0.2, 4], [0.25, 0.2, 0.2, 4], [0.25, 0.2, 0.2, 5]]
for val in hyperparams:
    WGRMF += [WGRMFDTIClassifier(similarity=sim, similarityCodes=codes, k=30, lambda_l=val[0], lambda_d=val[1], lambda_t=val[2], p=val[3])]
    WGRMFAddLog += [AddSingleDTIClassifier(modelType="LR", otherclassifier=WGRMFDTIClassifier(similarity=sim, similarityCodes=codes, k=30, lambda_l=val[0], lambda_d=0, lambda_t=val[2], p=val[3]))]
    WGRMFAddRF += [AddSingleDTIClassifier(modelType="RF", otherclassifier=WGRMFDTIClassifier(similarity=sim, similarityCodes=codes, k=30, lambda_l=val[0], lambda_d=0, lambda_t=val[2], p=val[3]))]

# run the AUC calculations for submatrix problem
models=[WeightedProfile, WeightedNNLR, WeightedNNRF, RLSWNN, RLSWNNSingleLog, RLSWNNSingleRF, CMF, CMFAddLog, CMFAddRF, WGRMF, WGRMFAddLog, WGRMFAddRF, RFOne, RFOneAddLog, RFOneAddRF]
plot1 = ([0, 1, 2], ["Weighted Profile", "Weighted NN + Log", "Weighted NN + RF"], outfolder + "WeightedNN")
plot2 = ([3, 4, 5], ["RLS-WNN", "RLS Target + Log", "RLS Target + RF"], outfolder + "RLS")
plot3 = ([6, 7, 8], ["CMF", "CMF + Log", "CMF + RF"], outfolder + "CMF")
plot4 = ([9, 10, 11], ["WGRMF", "WGRMF + Log", "WGRMF + RF"], outfolder + "WGRMF")
plot5 = ([12, 13, 14], ["RF One-Hot", "RF One-Hot + Log", "RF One-Hot + RF"], outfolder + "RFOneHot")
plotiVnfo = [plot1, plot2, plot3, plot4, plot5]
true_posmean, true_posstd, aucmean, aucstd, true_posutmean, true_posutstd, aucutmean, aucutstd, true_posudmean, true_posudstd, aucudmean, aucudstd, true_posudutmean, true_posudutstd, aucudutmean, aucudutstd, aucdiffmean, aucdiffstd, aucutdiffmean, aucutdiffstd, aucuddiffmean, aucuddiffstd, aucudutdiffmean, aucudutdiffstd, aucpertargets, true_pospertargetsall = AnalyzeDTIClassifier.SubmatrixROC(models, DTImatrix, codes, ligands, drugsim, repnum=1, plot=True, plotinfo=plotinfo,
hyperparams=True, splitFile=outfolder + "SplitInfo.npy", distanceFile=outfolder + "TargetDistance.npy", targetsim=sim)

# print out results
def printresults(index, modelname):
    with open(outfolder + modelname + "PerformanceSubmatrix.out", 'w') as f:
        f.write(str(aucmean[index]) + " " + str(aucstd[index]) + "\n")
        f.write(str(aucutmean[index]) + " " + str(aucutstd[index]) + "\n")
        f.write(str(aucudmean[index]) + " " + str(aucudstd[index]) + "\n")
        f.write(str(aucudutmean[index]) + " " + str(aucudutstd[index]) + "\n")
    np.savetxt(outfolder + modelname + "TruePos.out", true_posmean[index])
    np.savetxt(outfolder + modelname + "TruePosStd.out", true_posstd[index])
    np.savetxt(outfolder + modelname + "TruePosUT.out", true_posutmean[index])
    np.savetxt(outfolder + modelname + "TruePosUTStd.out", true_posutstd[index])
    np.savetxt(outfolder + modelname + "TruePosUD.out", true_posudmean[index])
    np.savetxt(outfolder + modelname + "TruePosUDStd.out", true_posudstd[index])
    np.savetxt(outfolder + modelname + "TruePosUDUT.out", true_posudutmean[index])
    np.savetxt(outfolder + modelname + "TruePosUDUTStd.out", true_posudutstd[index])
    np.savetxt(outfolder + modelname + "PerformancePerTarget.out", aucpertargets[:, index])

printresults(0, "WeightedProfile")
printresults(1, "WeightedNNLR")
printresults(2, "WeightedNNRF")
printresults(3, "RLSWNN")
printresults(4, "RLSWNNSingleLog")
printresults(5, "RLSWNNSingleRF")
printresults(6, "CMF")
printresults(7, "CMFAddLog")
printresults(8, "CMFAddRF")
printresults(9, "WGRMF")
printresults(10, "WGRMFAddLog")
printresults(11, "WGRMFAddRF")
printresults(12, "RFOne")
printresults(13, "RFOneAddLog")
printresults(14, "RFOneAddRF")

np.savetxt(outfolder + "AUCDiffMeanSubmatrix.out", aucdiffmean)
np.savetxt(outfolder + "AUCDiffStdSubmatrix.out", aucdiffstd)
np.savetxt(outfolder + "AUCUTDiffMeanSubmatrix.out", aucutdiffmean)
np.savetxt(outfolder + "AUCUTDiffStdSubmatrix.out", aucutdiffstd)
np.savetxt(outfolder + "AUCUDDiffMeanSubmatrix.out", aucuddiffmean)
np.savetxt(outfolder + "AUCUDDiffStdSubmatrix.out", aucuddiffstd)
np.savetxt(outfolder + "AUCUDUTDiffMeanSubmatrix.out", aucudutdiffmean)
np.savetxt(outfolder + "AUCUDUTDiffStdSubmatrix.out", aucudutdiffstd)
```
