## Supplementary material for "Using Single Protein/Ligand Binding Models to Predict Active Ligands for Unseen Proteins": Code and dataset: DTIModelsCompile.py

```
import numpy as np
import matplotlib
matplotlib.use('agg')
import matplotlib.pyplot as plt
import sys
import os
import subprocess
import math

# This script compiles the results from various runs into a single final set of plots. IN particular it takes averages and standard deviations of the AUCs and combines them for convenience and further analysis. You must specify the initial starting point, ending point, and the dataset. All results are stored in Analysis/DTI/<dataset>compiledrun<foldernum>.

dataset = sys.argv[1]
start = int(sys.argv[2])
end = int(sys.argv[3])
foldernum = int(sys.argv[4])

models = ["WeightedProfile", "WeightedNNLR", "WeightedNNRF", "RLSWNN", "RLSWNNSingleLog", "RLSWNNSingleRF", "CMF", "CMFAddLog", "CMFAddRF", "WGRMF", "WGRMFAddLog", "WGRMFAddRF", "RFOne", "RFOneAddLog", "RFOneAddRF"]
aucs = [[] for model in models]
true_posall = [np.array([]) for model in models]
aucuds = [[] for model in models]
true_posudall = [np.array([]) for model in models]
aucuts = [[] for model in models]
true_posutall = [np.array([]) for model in models]
aucuduts = [[] for model in models]
true_posudutall = [np.array([]) for model in models]
aucdiffs = [[[] for model in models] for model in models]
aucuddiffs = [[[] for model in models] for model in models]
aucutdiffs = [[[] for model in models] for model in models]
aucudutdiffs = [[[] for model in models] for model in models]

# read in all the data
for i in range(start, end):
    infolder = "Analysis/DTI/" + dataset + "run" + str(i) + "/"
    for j, model in enumerate(models):
        with open(infolder + model + "PerformanceSubmatrix.out", 'r') as f:
            aucs[j] += [float(f.readline().split(' ')[0])]
            aucuts[j] += [float(f.readline().split(' ')[0])]
            aucuds[j] += [float(f.readline().split(' ')[0])]
            aucuduts[j] += [float(f.readline().split(' ')[0])]
        if i == start:
            true_posall[j] = np.array([np.loadtxt(infolder + model + "TruePos.out")])
            true_posudall[j] = np.array([np.loadtxt(infolder + model + "TruePosUD.out")])
            true_posutall[j] = np.array([np.loadtxt(infolder + model + "TruePosUT.out")])
            true_posudutall[j] = np.array([np.loadtxt(infolder + model + "TruePosUDUT.out")])
        else:
            true_posall[j] = np.vstack((true_posall[j], np.loadtxt(infolder + model + "TruePos.out")))
            true_posudall[j] = np.vstack((true_posudall[j], np.loadtxt(infolder + model + "TruePosUD.out")))
            true_posutall[j] = np.vstack((true_posutall[j], np.loadtxt(infolder + model + "TruePosUT.out")))
            true_posudutall[j] = np.vstack((true_posudutall[j], np.loadtxt(infolder + model + "TruePosUDUT.out")))

for i in range(end-start):
    for k in range(len(models)):
        for l in range(len(models)):
            aucdiffs[k][l] += [aucs[k][i] - aucs[l][i]]
            aucutdiffs[k][l] += [aucuts[k][i] - aucuts[l][i]]
            aucuddiffs[k][l] += [aucuds[k][i] - aucuds[l][i]]
            aucudutdiffs[k][l] += [aucuduts[k][i] - aucuduts[l][i]]
    
aucs = np.array(aucs)
true_posall = np.array(true_posall)
aucuts = np.array(aucuts)
true_posutall = np.array(true_posutall)
aucuds = np.array(aucuds)
true_posudall = np.array(true_posudall)
aucuduts = np.array(aucuduts)
true_posudutall = np.array(true_posudutall)
aucdiffs = np.array(aucdiffs)
aucutdiffs = np.array(aucutdiffs)
aucuddiffs = np.array(aucuddiffs)
aucudutdiffs = np.array(aucudutdiffs)

print(np.shape(true_posall), np.shape(true_posutall), np.shape(true_posudall), np.shape(true_posudutall))

# compute means and standard deviations
aucmean = np.mean(aucs, axis=1)
aucstd = np.std(aucs, axis=1)/math.sqrt(end-start)
true_posmean = np.mean(true_posall, axis=1)
true_posstd = np.std(true_posall, axis=1)/math.sqrt(end-start)
aucudmean = np.mean(aucuds, axis=1)
aucudstd = np.std(aucuds, axis=1)/math.sqrt(end-start)
true_posudmean = np.mean(true_posudall, axis=1)
true_posudstd = np.std(true_posudall, axis=1)/math.sqrt(end-start)
aucutmean = np.mean(aucuts, axis=1)
aucutstd = np.std(aucuts, axis=1)/math.sqrt(end-start)
true_posutmean = np.mean(true_posutall, axis=1)
true_posutstd = np.std(true_posutall, axis=1)/math.sqrt(end-start)
aucudutmean = np.mean(aucuduts, axis=1)
aucudutstd = np.std(aucuduts, axis=1)/math.sqrt(end-start)
true_posudutmean = np.mean(true_posudutall, axis=1)
true_posudutstd = np.std(true_posudutall, axis=1)/math.sqrt(end-start)
aucdiffmean = np.mean(aucdiffs, axis=2)
aucdiffstd = np.std(aucdiffs, axis=2)/math.sqrt(end-start)
aucutdiffmean = np.mean(aucutdiffs, axis=2)
aucutdiffstd = np.std(aucutdiffs, axis=2)/math.sqrt(end-start)
aucuddiffmean = np.mean(aucuddiffs, axis=2)
aucuddiffstd = np.std(aucuddiffs, axis=2)/math.sqrt(end-start)
aucudutdiffmean = np.mean(aucudutdiffs, axis=2)
aucudutdiffstd = np.std(aucudutdiffs, axis=2)/math.sqrt(end-start)

print(np.shape(true_posmean))

# print out all the information
outfolder = "Analysis/DTI/" + dataset + "compiled" + str(foldernum) + "/"
subprocess.call("mkdir " + outfolder, shell=True)
subprocess.call("rm " + outfolder + "*", shell=True)

def printresults(index, modelname):
    with open(outfolder + modelname + "PerformanceSubmatrix.out", 'w') as f:
        f.write(str(aucmean[index]) + " " + str(aucstd[index]) + "\n")
        f.write(str(aucutmean[index]) + " " + str(aucutstd[index]) + "\n")
        f.write(str(aucudmean[index]) + " " + str(aucudstd[index]) + "\n")
        f.write(str(aucudutmean[index]) + " " + str(aucudutstd[index]) + "\n")
    np.savetxt(outfolder + modelname + "TruePos.out", true_posmean[index])
    np.savetxt(outfolder + modelname + "TruePosStd.out", true_posstd[index])
    np.savetxt(outfolder + modelname + "TruePosUT.out", true_posutmean[index])
    np.savetxt(outfolder + modelname + "TruePosUTStd.out", true_posutstd[index])
    np.savetxt(outfolder + modelname + "TruePosUD.out", true_posudmean[index])
    np.savetxt(outfolder + modelname + "TruePosUDStd.out", true_posudstd[index])
    np.savetxt(outfolder + modelname + "TruePosUDUT.out", true_posudutmean[index])
    np.savetxt(outfolder + modelname + "TruePosUDUTStd.out", true_posudutstd[index])

for i, model in enumerate(models):
    printresults(i, model)

# make plots
plot0 = ([0, 1, 2], ["Weighted Profile", "Weighted NN + Log", "Weighted NN + RF"], outfolder + "WeightedNN")
plot1 = ([3, 4, 5], ["RF One-Hot", "RF One-Hot + Log", "RF One-Hot + RF"], outfolder + "RFOneHot")
plot2 = ([6, 7, 8], ["RLS-WNN", "RLS Target + Log", "RLS Target + RF"], outfolder + "RLS")
plot3 = ([9, 10, 11], ["CMF", "CMF + Log", "CMF + RF"], outfolder + "CMF")
plot4 = ([12, 13, 14], ["WGRMF", "WGRMF + Log", "WGRMF + RF"], outfolder + "WGRMF")
plotinfo = [plot0, plot1, plot2, plot3, plot4]

mean_fpr = np.linspace(0, 1, 1000)
for plotmodels, plotlabels, plotfiles in plotinfo:
    fig, ax = plt.subplots()
    for i, k in enumerate(plotmodels):
        ax.errorbar(mean_fpr, true_posmean[k], yerr=true_posstd[k], label=plotlabels[i])
    ax.set_xlim([0, 1])
    ax.set_ylim([0, 1])
    ax.set_xlabel("False positive")
    ax.set_ylabel("True positive")
    plt.legend()
    plt.savefig(plotfiles + "AllROC.jpg")

    fig, ax = plt.subplots()
    for i, k in enumerate(plotmodels):
        ax.errorbar(mean_fpr, true_posutmean[k], yerr=true_posutstd[k], label=plotlabels[i])
    ax.set_xlim([0, 1])
    ax.set_ylim([0, 1])
    ax.set_xlabel("False positive")
    ax.set_ylabel("True positive")
    plt.legend()
    plt.savefig(plotfiles + "UnknownTargetROC.jpg")

    fig, ax = plt.subplots()
    for i, k in enumerate(plotmodels):
        ax.errorbar(mean_fpr, true_posudmean[k], yerr=true_posudstd[k], label=plotlabels[i])
    ax.set_xlim([0, 1])
    ax.set_ylim([0, 1])
    ax.set_xlabel("False positive")
    ax.set_ylabel("True positive")
    plt.legend()
    plt.savefig(plotfiles + "UnknownDrugROC.jpg")
    
    fig, ax = plt.subplots()
    for i, k in enumerate(plotmodels):
        ax.errorbar(mean_fpr, true_posudutmean[k],yerr=true_posudutstd[k], label=plotlabels[i])
    ax.set_xlim([0, 1])
    ax.set_ylim([0, 1])
    ax.set_xlabel("False positive")
    ax.set_ylabel("True positive")
    plt.legend()
    plt.savefig(plotfiles + "BothUnknownROC.jpg")
```
