## Supplementary material for "Using Single Protein/Ligand Binding Models to Predict Active Ligands for Unseen Proteins": Code and dataset: OneHotDTIClassifier.py

```
import numpy as np
from sklearn.linear_model import LogisticRegression
from sklearn.ensemble import RandomForestClassifier
import sys
import pickle

# This runs a simple classifier (either LR or RF at present) on the one-hot fingerprints of everything combined.

class OneHotDTIClassifier:

    def __init__(self, sequence, model, hyperparams):
        self.sequence = sequence
        self.model = model
        self.hyperparams = hyperparams

    # submatrix method: takes in a training matrix, ligand list, list of training/test proteins and ligands, and outputs a completed test matrix
    def fit_predict_submatrix(self, trainmat, trainmatlog, trainmatrf, ligands, codes, trainprotvals, testprotvals, trainligvals, testligvals, X_trainonehotseq, X_testonehotseq, drugsim):
        trainmat = np.copy(trainmat)

        if self.sequence == 'Seq':
            X_trainonehotseq = X_trainonehotseq.tolil()
            X_testonehotseq = X_testonehotseq.tolil()
            X_train = X_trainonehotseq[:,:-1]
            y_train = X_trainonehotseq[:,-1].toarray().flatten()
            X_test = X_testonehotseq[:,2:]
            indices = X_testonehotseq[:,:2].toarray()
        else:
            raise NotImplementedError

        if self.model == 'LR':
            log = LogisticRegression(C=self.hyperparams[0], n_jobs=-1)
            log.fit(X_train, y_train)
            y_test = log.predict_proba(X_test)[:,1]
        elif self.model == 'RF':
            rf = RandomForestClassifier(n_estimators=self.hyperparams[0], max_depth=self.hyperparams[1], n_jobs=-1)
            rf.fit(X_train, y_train)
            y_test = rf.predict_proba(X_test)[:,1]
        else:
            raise NotImplementedError

        for index, val in enumerate(y_test):
            trainmat[int(indices[index,0])][int(indices[index,1])] = val

        return trainmat
```
