## Supplementary material for "Using Single Protein/Ligand Binding Models to Predict Active Ligands for Unseen Proteins": Code and dataset: OneHotSingleDTIClassifier.py

```
import numpy as np
from sklearn.linear_model import LogisticRegression
from sklearn.ensemble import RandomForestClassifier, RandomForestRegressor
import sys
from scipy import sparse
import pickle

# This improves on the one-hot DTI classifier by using a single protein/many ligand model initially for the known protein/unknown ligand problem. For RF, we incorporate the initial predictions into the final result and secretly use a regressor instead.

class OneHotSingleDTIClassifier:

    def __init__(self, sequence, model, hyperparams, singlemodel):
        self.sequence = sequence
        self.model = model
        self.hyperparams = hyperparams
        self.singlemodel = singlemodel

    # submatrix method: takes in a training matrix, ligand list, list of training/test proteins and ligands, and outputs a completed test matrix
    def fit_predict_submatrix(self, trainmat, trainmatlog, trainmatrf, ligands, codes, trainprotvals, testprotvals, trainligvals, testligvals, X_trainonehotseq, X_testonehotseq, drugsim):
        if self.singlemodel == 'LR':
            trainmat = np.copy(trainmatlog)
        elif self.singlemodel == 'RF':
            trainmat = np.copy(trainmatrf)
        else:
            raise NotImplementedError
       
        # get the features
        if self.sequence == 'Seq':
            X_trainonehotseq = X_trainonehotseq.tolil()
            X_testonehotseq = X_testonehotseq.tolil()
            X_train = X_trainonehotseq[:,:-1]
            y_train = X_trainonehotseq[:,-1].toarray().flatten()
            X_test = X_testonehotseq[:,2:]
            indices = X_testonehotseq[:,:2].toarray()
        else:
            raise NotImplementedError

        # run the appropriate model
        if self.model == 'LR':
            log = LogisticRegression(C=self.hyperparams[0], n_jobs=-1)
            log.fit(X_train, y_train)
            y_test = log.predict_proba(X_test)[:,1]
        elif self.model == 'RF':
            # add in the values generated by the single protein/many ligand model
            mask = np.ones(X_test.shape[0], dtype=bool)
            for j, row in enumerate(indices):
                if row[0] in trainprotvals:
                    mask[j] = False
                    X_train = sparse.vstack((X_train, X_test[j].toarray()))
                    y_train = np.append(y_train, trainmat[int(row[0])][int(row[1])])
            X_test = X_test.tocsr()
            X_test = X_test[mask]
            indices = indices[mask]
            
            rf = RandomForestRegressor(n_estimators=self.hyperparams[0], max_depth=self.hyperparams[1], n_jobs=-1, max_features=1/3)
            rf.fit(X_train, y_train)
            y_test = rf.predict(X_test)
        else:
            raise NotImplementedError
        
        # fill in, but only for the test proteins
        for index, val in enumerate(y_test):
            if indices[index, 0] in testprotvals:
                trainmat[int(indices[index,0])][int(indices[index,1])] = val

        return trainmat
```
