## Supplementary material for "Using Single Protein/Ligand Binding Models to Predict Active Ligands for Unseen Proteins": Code and dataset: ParseCHEMBLAll.py

```
import numpy as np
from chembl_webresource_client.new_client import new_client as nc
from rdkit import Chem
from rdkit.Chem import AllChem

# This script runs through all ChEMBL target ids and provides output files with the active CHEMBL codes and the fingerprints of the active ligands. Non-active ligands are stored separately with fingerprint data. Activities are all stored.
# ChEMBL_Ligand_Data/ has the ChEMBL codes, Active_Fingerprint_Data/ has the fingerprints and activities of active ligands (activity in uM). Other_Fingerprint_Data/ has the fingerprints and activities of other ligands (activity in uM).

# computes Morgan fingerprint of the molecule in activity and adds it to actives, also adding the activity
def toMorgan(activity, actives):
    # now get the SMILES and go to rdkit
    try:
        smiles = activity['canonical_smiles']
        value = float(activity['standard_value'])
        units = activity['standard_units']
        if smiles is not None and Chem.MolFromSmiles(smiles) is not None:
            mol = Chem.MolFromSmiles(smiles)
        
            # get and use Morgan 3 fingerprint
            Morgan3 = AllChem.GetMorganFingerprintAsBitVect(mol, 3).ToBitString()
            Morgan3arr = np.array(list(map(int, Morgan3)))
            if units == "nM":
                value = value/1000
            Morgan3arr = np.append(Morgan3arr, value)
            if len(actives) == 0:
                actives = Morgan3arr
            else:
                actives = np.vstack((actives, Morgan3arr))
        
        else:
            print(smiles)
    except KeyError as error:
        print(error)
        pass
    return actives


# get all the targets
targetlist = nc.target.filter()
for target in targetlist:
    # use CHEMBL to find the activities
    testtarget = target['target_chembl_id']
    print(testtarget)
    activities = nc.activity.filter(target_chembl_id__in=testtarget).only(['target_chembl_id', 'molecule_chembl_id', 'standard_type', 'standard_units', 'standard_value', 'canonical_smiles'])

    # search through them looking for actives. Active criteria: some binding constant < 1 uM
    outFile = open("ChEMBL_Ligand_Data/" + testtarget + "Ligands.txt", "w")
    valid = ["Ki", "Kd", "IC50", "EC50", "Kic", "Kieq", "KiH", "Log Ki", "logKi", "logEC50", "Potency", "KiL"]
    ligands = set()
    otherL = set()
    actives = np.array([])
    others = np.array([])
    for activity in activities:
        if activity['standard_type'] in valid and activity['standard_value'] is not None:
            value = float(activity['standard_value'])
            units = activity['standard_units']
            mol = activity['molecule_chembl_id']
    
            # is it active?
            if ((value < 1000 and units == "nM") or (value < 1 and units == "uM")) and (mol not in ligands and mol not in otherL):
                outFile.write(mol + "\n")
                ligands.add(mol)
                actives = toMorgan(activity, actives)

            # if not active, get the activity anyway
            elif mol not in ligands and mol not in otherL and ((value >= 1000 and units == "nM") or (value >= 1 and units == "uM")):
                otherL.add(mol)
                others = toMorgan(activity, others)
    
    print(np.shape(actives))
    np.savetxt('Active_Fingerprint_Data/' + testtarget, actives)
    np.savetxt('Other_Fingerprint_Data/' + testtarget, others)
```
