## Supplementary material for "Using Single Protein/Ligand Binding Models to Predict Active Ligands for Unseen Proteins": Code and dataset: PubChemDecoys.py

```
import numpy as np
from chembl_webresource_client.new_client import new_client as nc
from rdkit import Chem
from rdkit.Chem import AllChem
from bioservices import UniProt, EUtils
import time

# This script runs through all ChEMBL targets and identifies compounds in the PubChem database that have been labeled as an inactive on some assay for that target, to be used as decoys. Compounds that have been labeled as both inactive and active are stored separately.
# The pause times throughout the script are essential to avoid violating the terms of use for EUtils.
# Output files: Inactive SMILES to Inactive_SMILES/, inactive fingerprints to Inactive_Fingerprint_Data/, conflict SMILES to Conflict_SMILES/.

# chunk an array up for EUtils
def chunk(arr, n):
    for i in range(0, len(arr), n):
        yield arr[i:i+n]

# get all the targets
targetlist = nc.target.filter()
for target in targetlist:
    try:
        targetid = target['target_chembl_id']
        print(targetid)
        outFile = open("Inactive_SMILES/" + targetid + ".txt", "w")
    
        # get the uniprot ID
        uniprot = target['target_components'][0]['accession']
        s = EUtils()
        # get the protein ID
        search = s.ESearch("protein", uniprot)
    
        # get a list of all the associated assays
        link = s.ELink(dbfrom="protein", dbto="pcassay", id=search['idlist'], linkname="protein_pcassay_target")
        link = s.parse_xml(link, 'EUtilsParser')
        if 'LinkSet' in link.eLinkResult.keys() and 'LinkSetDb' in link.eLinkResult.LinkSet.keys():
            if len(link.eLinkResult.LinkSet.LinkSetDb.Link) == 1:
                assays = [link.eLinkResult.LinkSet.LinkSetDb.Link['Id']]
            else:
                assays = [assay['Id'] for assay in link.eLinkResult.LinkSet.LinkSetDb.Link]
            time.sleep(1)
    
            # get all the active and inactive compounds
            inactives = set()
            actives = set()
            for part in chunk(assays, 200):
                # get inactives via ELink
                ret = s.ELink("pcassay", "pccompound", id=part, linkname="pcassay_pccompound_inactive")
                ret = s.parse_xml(ret, 'EUtilsParser')
                if 'LinkSetDb' in ret.eLinkResult.LinkSet.keys():
                    if len(ret.eLinkResult.LinkSet.LinkSetDb.Link) == 1:
                        inactive = [ret.eLinkResult.LinkSet.LinkSetDb.Link['Id']]
                    else:
                        inactive = [compound['Id'] for compound in ret.eLinkResult.LinkSet.LinkSetDb.Link]
                inactives.update(inactive)
        
                # get actives via ELink
                ret = s.ELink("pcassay", "pccompound", id=part, linkname="pcassay_pccompound_active")
                ret = s.parse_xml(ret, 'EUtilsParser')
                if 'LinkSetDb' in ret.eLinkResult.LinkSet.keys():
                    if len(ret.eLinkResult.LinkSet.LinkSetDb.Link) == 1:
                        active = [ret.eLinkResult.LinkSet.LinkSetDb.Link['Id']]
                    else:
                        active = [compound['Id'] for compound in ret.eLinkResult.LinkSet.LinkSetDb.Link]
                actives.update(active)
        
                time.sleep(1)
            
            # get the conflicts
            conflicts = actives.intersection(inactives)
            inactives = inactives - conflicts
            inactiveArr = np.array([])
        
            # get SMILES from inactives
            for part in chunk(list(inactives), 200):
                # get smiles
                ret = s.ESummary("pccompound", id=part)
                for val in part:
                    smiles = ret[val]['canonicalsmiles']
                    outFile.write(smiles + "\n")
                    # convert to fingerprint
                    try:
                        if smiles is not None and Chem.MolFromSmiles(smiles) is not None:
                            mol = Chem.MolFromSmiles(smiles)
                
                            # get and use Morgan 3 fingerprint
                            Morgan3 = AllChem.GetMorganFingerprintAsBitVect(mol, 3).ToBitString()
                            Morgan3arr = np.array(list(map(int, Morgan3)))
                            if len(inactiveArr) == 0:
                                inactiveArr = Morgan3arr
                            else:
                                inactiveArr = np.vstack((inactiveArr, Morgan3arr))
                
                        else:
                            print(smiles)
                    except KeyError as error:
                        print(error)
                        pass
                time.sleep(1)
        
            np.savetxt('Inactive_Fingerprint_Data/' + targetid, inactiveArr)
            outFile.close()
        
            # get SMILES from conflicts
            outFile = open("Conflict_SMILES/" + targetid + ".txt", "w")
            for part in chunk(list(conflicts), 200):
                ret = s.ESummary("pccompound", id=part)
                for val in part:
                    smiles = ret[val]['canonicalsmiles']
                    outFile.write(smiles + "\n")
                time.sleep(1)
        outFile.close()

    except (TypeError, KeyError) as error:
        print(error)
        pass
```
