## Supplementary material for "Using Single Protein/Ligand Binding Models to Predict Active Ligands for Unseen Proteins": Code and dataset: README

There are 6 scripts provided to recreate the results of this paper and also generate a database of protein/ligand binding data that may prove useful for any other data analysis.

1. ParseCHEMBLAll.py - parses the ChEMBL database and finds all active ligands for every protein target stored in the database. Stores Morgan 3 fingerprints, activities, and activities for all inactive ligands just in case.

2. PubChemDecoys.py - parses the PubChem database for inactives as described in the paper. Stores all inactive fingerprints along with a small number of conflicts (ligands that were both inactive and active).

3. SequenceListCompile.py - generates the list of sequences from UniProt and computes a sequence similarity matrix using pHMMER. Stores both the sequence list and the sequence similarity matrices.

4. CompileLigandList.py - generates the fingerprints for all the ligands, along with the fingerprint similarity matrices and the one-hot fingerprint features used in the one-hot model. Stores all of this information for use during the classifier runs.

5. DTIModelsCompare.py - main script to run and analyze results from all the DTI models. Calls a number of the other scripts, which can be modified as necessary but should not need modification or running to reproduce the results of the paper. This can/should be run in a multithreaded/distributed fashion on a single core, and what operations can be easily distributed are by using numpy or sklearn functions.

6. DTIModelsCompile.py - combines results from multiple runs into averages and standard deviations, which can then be processed for further analysis.

The other helper classes provided (in particular AnalyzeDTIClassifier.py) help set up the problem and evaluate the various models. The model classes can be used as models for building further models for this dataset.

You will need to download all the ChEMBL molecules from the database directly into a file chembl_<version>_chemreps.txt and change the version number in AnalyzeDTIClassifier.py as appropriate.

Packages you will need:
numpy
rdkit
chembl_webresource_client
bioservices
BioPDB
HMMER (not a package - just the phmmer command line)
urllib
scipy
sklearn
matplotlib
