## Supplementary material for "Using Single Protein/Ligand Binding Models to Predict Active Ligands for Unseen Proteins": Code and dataset: RLSWNNDTIClassifier.py

```
import numpy as np
import math
from numpy import linalg
from scipy.spatial.distance import pdist, squareform
from scipy.sparse.linalg import eigsh
import sys
import pickle

# This class implements the RLS-WNN classifier as described in van Laarhoven 2013 (with the WNN preprocessing for improvement performance). It runs weighted-nearest-neighbor for preprocessing and then uses the RLS-kron formulation with the Gaussian interaction profile incorporated. It also uses eigenvalues to speed up inversion of the matrix and avoid constructing it directly; due to runtime issues, we only take the top k eigenvalues of the drug similarity matrix. eta is the decoy used in the WNN calculation, alpha determines how much influence the GIP kernel has on the algorithm, and sigma is a regularization parameter on RLS.

class RLSWNNDTIClassifier:

    def __init__(self, similarity, similarityCodes, eta, alpha, sigma, k):
        self.similarity = similarity
        self.similarityCodes = similarityCodes
        self.eta = eta
        self.alpha = alpha
        self.sigma = sigma
        self.k = k

    # submatrix method: takes in a training matrix, ligand list, list of training/test proteins and ligands, and outputs a completed test matrix
    def fit_predict_submatrix(self, trainmat, trainmatlog, trainmatrf, ligands, codes, trainprotvals, testprotvals, trainligvals, testligvals, X_trainonehotseq, X_testonehotseq, drugsim):
        trainmat = np.copy(trainmat)
        trainmat[trainmat == -1] = 0
        targetsim = np.zeros((len(codes), len(codes)))
        for i, code in enumerate(codes):
            for j, othercode in enumerate(codes):
                targetsim[i][j] = self.similarity[self.similarityCodes.index(code)][self.similarityCodes.index(othercode)]

        # WNN preprocessing for both drugs and targets
        for i in range(np.shape(ligands)[0]):
            if (trainmat[:,i] == 0).all():
                indices = np.flip(np.argsort(drugsim[i,:]))[1:50]
                powers = np.array([math.pow(self.eta, j) for j in range(49)])
                trainmat[:,i] = np.dot(powers, trainmat[:,indices].T).T
        for i in range(len(codes)):
            if (trainmat[i,:] == 0).all():
                indices = np.flip(np.argsort(targetsim[i,:]))[1:]
                powers = np.array([math.pow(self.eta, j) for j in range(len(codes)-1)])
                trainmat[i,:] = np.dot(powers, trainmat[indices,:])

        # update the kernel matrices with the GIP kernel
        dist = squareform(pdist(trainmat, metric='euclidean'))
        gamma = np.sum(trainmat)/np.shape(trainmat)[0]
        kernel = np.exp(-dist/gamma)
        targetsim = self.alpha*targetsim + (1-self.alpha)*kernel

        dist = squareform(pdist(trainmat.T, metric='euclidean'))
        gamma = np.sum(trainmat)/np.shape(trainmat)[1]
        kernel = np.exp(-dist/gamma)
        drugsim = self.alpha*drugsim + (1-self.alpha)*kernel

        # now run the RLS algorithm with the eigenvalue formulation
        la, va = linalg.eig(targetsim)
        lb, vb = eigsh(drugsim, k=self.k)
        sys.stdout.flush()
        l = np.outer(la, np.array([lb]).T)
        l = l/(l + self.sigma)
        m1 = np.dot(np.dot(va.T, trainmat), vb)
        m2 = m1*l
        trainmat = np.dot(np.dot(va, m2), vb.T)

        return trainmat
```
