## Supplementary material for "Using Single Protein/Ligand Binding Models to Predict Active Ligands for Unseen Proteins": Code and dataset: RLSWNNSingleDTIClassifier.py

```
import numpy as np
import math
from numpy import linalg
from scipy.spatial.distance import pdist, squareform
from scipy.sparse.linalg import eigsh
import sys
from sklearn.linear_model import LogisticRegression
from sklearn.ensemble import RandomForestClassifier
import pickle

# This class implements the RLS-WNN classifier as described in van Laarhoven 2013 (with the WNN preprocessing for improvement performance) with using a single protein/many ligand model as preprocessing. It runs weighted-nearest-neighbor and the appropriate model for preprocessing and then uses the RLS formulation with the Gaussian interaction profile incorporated. It only runs the RLS formulation on the targets, since that is all that is necessary, and therefore omits the eigenvalue formulation. eta is the decoy used in the WNN calculation, alpha determines how much influence the GIP kernel has on the algorithm, and sigma is a regularization parameter on RLS.

class RLSWNNSingleDTIClassifier:

    def __init__(self, similarity, similarityCodes, eta, alpha, sigma, modelType):
        self.similarity = similarity
        self.similarityCodes = similarityCodes
        self.eta = eta
        self.alpha = alpha
        self.sigma = sigma
        self.modelType = modelType

    # submatrix method: takes in a training matrix, ligand list, list of training/test proteins and ligands, and outputs a completed test matrix
    def fit_predict_submatrix(self, trainmat, trainmatlog, trainmatrf, ligands, codes, trainprotvals, testprotvals, trainligvals, testligvals, X_trainonehotseq, X_testonehotseq, drugsim):
        targetsim = np.zeros((len(codes), len(codes)))
        for i, code in enumerate(codes):
            for j, othercode in enumerate(codes):
                targetsim[i][j] = self.similarity[self.similarityCodes.index(code)][self.similarityCodes.index(othercode)]
        if self.modelType == 'LR':
            trainmat = np.copy(trainmatlog)
        elif self.modelType == 'RF':
            trainmat = np.copy(trainmatrf)
        else:
            raise NotImplementedError
        trainmat[trainmat == -1] = 0

        # WNN preprocessing for targets
        for i in range(len(codes)):
            if (trainmat[i,:] == 0).all():
                indices = np.flip(np.argsort(targetsim[i,:]))[1:]
                powers = np.array([math.pow(self.eta, j) for j in range(len(codes)-1)])
                trainmat[i,:] = np.dot(powers, trainmat[indices,:])

        # update the kernel matrices with the GIP kernel
        dist = squareform(pdist(trainmat, metric='euclidean'))
        gamma = np.sum(trainmat)/np.shape(trainmat)[0]
        kernel = np.exp(-dist/gamma)
        targetsim = self.alpha*targetsim + (1-self.alpha)*kernel

        # now run the RLS algorithm
        inv = np.linalg.inv(targetsim + self.sigma*np.identity(np.shape(targetsim)[0]))
        trainmat = np.dot(np.dot(targetsim, inv), trainmat)

        return trainmat
```
