## Supplementary material for "Using Single Protein/Ligand Binding Models to Predict Active Ligands for Unseen Proteins": Code and dataset: SequenceListCompile.py

```
from chembl_webresource_client.new_client import new_client as nc
from bioservices import UniProt
import sys
import __main__
import os
import Bio.PDB as pdb
import urllib.request as urllib
import json
import numpy as np
import math

# This script takes in a dataset, consisting of a code list for all the proteins, and retrieves the relevant sequences from UniProt, accessed by ChEMBL ID. It then runs a sequence similarity computation using pHMMER for use in the DTI code.
# The sequences are stored in Real_Sequences/ by ChEMBL ID and the similarity matrices are stored in Similarity/ by the dataset name. A number of temporary files are generated which can be deleted once the code is run.

dataset = sys.argv[1]
codes = []
with open('codes_' + dataset + '.txt', 'r') as f:
    for line in f:
        codes += [line[:-1]]

sequences = []
for code in codes:
    # get UniProt ID
    target = nc.target
    res = target.get('CHEMBL' + code)
    uniprot = res['target_components'][0]['accession']

    # get sequence from UniProt
    u = UniProt()
    sequence = u.get_fasta_sequence(uniprot)
    with open("Real_Sequences/CHEMBL" + code, 'w') as f:
        f.write('>CHEMBL' + code + '\n')
        f.write(sequence)
        sequences += [sequence]

sim = np.identity(len(codes))
for i in range(len(codes)):
    with open("tempfile.fa", 'w') as f:
        f.write(">CHEMBL" + codes[i] + "\n")
        f.write(sequences[i] + "\n")
    with open("tempdatabase.fa", 'w') as f:
        for j in range(len(codes)):
            f.write(">CHEMBL" + codes[j] + "\n")
            f.write(sequences[j] + "\n")
    os.system("phmmer -o throwaway.out --tblout temp.out -T 0 --F1 1 --F2 1 --F3 1 tempfile.fa tempdatabase.fa")
    with open("temp.out", 'r') as f:
        for line in f:
            if line[0] == "#":
                continue
            split = line.split(" ")
            split = list(filter(None, split))
            othercode = split[0]
            simval = float(split[5])
            index = codes.index(othercode[6:])
            sim[i][index] = max(sim[i][index], simval)
            sim[index][i] = max(sim[index][i], simval)

for i in range(len(codes)):
    for j in range(i+1, len(codes)):
        sim[i][j] = sim[i][j]/math.sqrt(sim[i][i]*sim[j][j])
        sim[j][i] = sim[j][i]/math.sqrt(sim[i][i]*sim[j][j])
        if sim[i][j] == 0:
            sim[i][j] = 1e-6
            sim[j][i] = 1e-6
for i in range(len(codes)):
    sim[i][i] = 1

minval = np.linalg.eigh(sim)[0][0]
if minval < 0:
    temp = sim + (-minval+0.001)*np.identity(np.shape(sim)[0])
    sim = temp/(1-minval+0.001)

np.savetxt('Similarity/Similarity_Sequence_' + dataset + '.out', sim)
```
