## Supplementary material for "Using Single Protein/Ligand Binding Models to Predict Active Ligands for Unseen Proteins": Code and dataset: WeightedNNDTIClassifier.py

```
import numpy as np
from sklearn.externals import joblib
from sklearn.linear_model import LogisticRegression
from sklearn.ensemble import RandomForestClassifier
import math

# This class implements a weighted NN based classifier for the DTI problem. It extrapolates a ligand profile for a given protein using the given model type and the stored model, and then uses  the given similarity matrix to compute the predictions for the zero-learning case. For the submatrix case, it proceeds similarly.

class WeightedNNDTIClassifier:

    def __init__(self, similarity, similarityCodes, modelType="LR"):
        self.similarity = similarity
        self.similarityCodes = similarityCodes
        self.modelType = modelType

    # submatrix method: takes in a training matrix, ligand list, list of training/test proteins and ligands, and outputs a completed test matrix
    def fit_predict_submatrix(self, trainmat, trainmatlog, trainmatrf, ligands, codes, trainprotvals, testprotvals, trainligvals, testligvals, X_trainonehotseq, X_testonehotseq, drugsim):
        if self.modelType == 'LR':
            trainmat = np.copy(trainmatlog)
        elif self.modelType == 'RF':
            trainmat = np.copy(trainmatrf)
        else:
            raise NotImplementedError

        # now take a look at the unknowns
        for i in testprotvals:
            code = codes[i]
            # find closest k training targets
            similarities = np.zeros((len(codes)))
            for j in trainprotvals:
                targetcode = codes[j]
                similarities[j] = self.similarity[self.similarityCodes.index(code)][self.similarityCodes.index(targetcode)]
            
            trainmat[i,:] = np.dot(similarities, trainmat)/np.sum(similarities)
        
        return trainmat
```
