## Supplementary material for "Using Single Protein/Ligand Binding Models to Predict Active Ligands for Unseen Proteins": Code and dataset: WeightedProfileDTIClassifier.py

```
import numpy as np
from scipy.spatial.distance import pdist, squareform

# This implements the weighted profile classifier from Yamanishi et al. (2008). It just picks the closest neighbor and evaluates the model based on that. Target similarity is whatever is given; drug similarity is Tanimoto similarity.

class WeightedProfileDTIClassifier:

    def __init__(self, similarity, similarityCodes):
        self.similarity = similarity
        self.similarityCodes = similarityCodes

    # submatrix method: takes in a training matrix, ligand list, list of training/test proteins and ligands, and outputs a completed test matrix
    def fit_predict_submatrix(self, trainmat, trainmatlog, trainmatrf, ligands, codes, trainprotvals, testprotvals, trainligvals, testligvals, X_trainonehotseq, X_testonehotseq, drugsim):
        trainmat = np.copy(trainmat)
        # set the unknowns to 0
        trainmat[trainmat==-1] = 0

        # grab the drug-weighted measurement
        trainmatdrug = (np.dot(drugsim, trainmat.T)/(np.sum(drugsim, axis=0)[:,None])).T
        trainmatdrug[trainmat==1] = 1

        # grab the measurement from the closest target
        targetsim = np.zeros((len(codes), len(codes)))
        for i, code in enumerate(codes):
            for j, othercode in enumerate(codes):
                targetsim[i][j] = self.similarity[self.similarityCodes.index(code)][self.similarityCodes.index(othercode)]
        trainmattarget = np.dot(targetsim, trainmat)/(np.sum(targetsim, axis=0)[:,None])
        trainmattarget[trainmat==1] = 1

        return (trainmatdrug + trainmattarget)/2
```
