## Supplementary material for "Using Single Protein/Ligand Binding Models to Predict Active Ligands for Unseen Proteins": Code and dataset: WGRMFDTIClassifier.py

```
import numpy as np
from numpy import linalg
from scipy.sparse.linalg import svds
from scipy import sparse
import pickle

# This class implements the WGRMF classifier as described in Ezzat 2017. It attempts to factor the drug/target interaction matrix trainmat as AB^T with regularization penalties lambda_l (overall matrix norm), lambda_d (deviation from drug similarity as measured by the normalized Laplacian), and lambda_t (ditto for target similarity). k is the number of latent features (must be < than the number of targets), and p is the number of nearest neighbors left after preprocessing.

class WGRMFDTIClassifier:

    def __init__(self, similarity, similarityCodes, k, lambda_l, lambda_d, lambda_t, p):
        self.similarity = similarity
        self.similarityCodes = similarityCodes
        self.k = k
        self.lambda_l = lambda_l
        self.lambda_d = lambda_d
        self.lambda_t = lambda_t
        self.p = p

    # submatrix method: takes in a training matrix, ligand list, list of training/test proteins and ligands, and outputs a completed test matrix
    def fit_predict_submatrix(self, trainmat, trainmatlog, trainmatrf, ligands, codes, trainprotvals, testprotvals, trainligvals, testligvals, X_trainonehotseq, X_testonehotseq, drugsim):
        self.k = min(self.k, np.shape(trainmat)[0]-1)
        trainmat = np.copy(trainmat)
        W = (trainmat != -1)
        trainmat[trainmat == -1] = 0
        targetsim = np.zeros((len(codes), len(codes)))
        for i, code in enumerate(codes):
            for j, othercode in enumerate(codes):
                targetsim[i][j] = self.similarity[self.similarityCodes.index(code)][self.similarityCodes.index(othercode)]
        losses = []

        # sparsify drug and target similarity matrices
        drugsim = np.copy(drugsim)
        for i in range(len(codes)):
            indices = np.argsort(targetsim[i,:])
            targetsim[i, indices[:-self.p-1]] = 0
        targetsim = (targetsim + targetsim.T)/2
        for i in range(np.shape(ligands)[0]):
            indices = np.argsort(drugsim[i,:])
            drugsim[i, indices[:-self.p-1]] = 0
        drugsim = (drugsim + drugsim.T)/2

        # compute Laplacian matrices
        sumd = np.sum(drugsim, axis=0)
        Dd = np.diag(sumd)
        powerd = np.power(sumd, -0.5)
        Ld = Dd - drugsim
        Ld = np.array([powerd]).T*Ld*powerd
        sumt = np.sum(targetsim, axis=0)
        Dt = np.diag(sumt)
        powert = np.power(sumt, -0.5)
        Lt = Dt - targetsim
        Lt = np.array([powert]).T*Lt*powert

        # initialize latent feature matrices and weight matrix
        trainmatsparse = sparse.lil_matrix(trainmat)
        u, s, vh = svds(trainmatsparse, k=self.k)
        v = vh.T
        A = u*np.sqrt(s)
        B = v*np.sqrt(s)
        loss = np.linalg.norm(np.multiply(W, trainmat - np.dot(A, B.T))) + self.lambda_l*(np.linalg.norm(A) + np.linalg.norm(B)) + self.lambda_d*np.trace(np.dot(np.dot(B.T, Ld), B)) + self.lambda_t*np.trace(np.dot(np.dot(A.T, Lt), A))
        losses += [loss]

        # run the iterations
        lambda_d_Ld = self.lambda_d*Ld
        lambda_t_Lt = self.lambda_t*Lt
        lambda_l_ik = self.lambda_l*np.identity(self.k)
        H = np.multiply(W, trainmat)

        # see the Ezzat paper's Github for the code I reimplemented here
        notdone = True
        count = 0
        while notdone and count < 20:
            count += 1
            A_old = A
            B_old = B
            numA = np.dot(H, B) - np.dot(lambda_t_Lt, A_old)
            for a in range(np.shape(A)[0]):
                A[a,:] = linalg.solve((np.dot(B.T*W[a,:], B) + lambda_l_ik).T, numA[a,:].T).T
            numB = np.dot(H.T, A) - np.dot(lambda_d_Ld, B_old)
            for b in range(np.shape(B)[0]):
                B[b,:] = linalg.solve((np.dot(A.T*W[:,b], A) + lambda_l_ik).T, numB[b,:].T).T
            loss = np.linalg.norm(np.multiply(W, trainmat - np.dot(A, B.T))) + self.lambda_l*(np.linalg.norm(A) + np.linalg.norm(B)) + self.lambda_d*np.trace(np.dot(np.dot(B.T, Ld), B)) + self.lambda_t*np.trace(np.dot(np.dot(A.T, Lt), A))
            losses += [loss]
            if losses[-1] > losses[-2] - 1:
                notdone = False

        # return the answer
        return np.dot(A, B.T)
```
